# Parameterization of the PA endonuclease bimetallic center reveals the dynamics of clinically relevant mutations

**DOI:** 10.64898/2026.06.12.731895

**Authors:** Luxuan Wang, Pengfei Li, Terra Sztain

**Affiliations:** Department of Medicinal Chemistry, University of Michigan, Ann Arbor, MI, USA; Department of Chemistry and Biochemistry, Loyola University Chicago, Chicago, IL, USA; Department of Biophysics, University of Michigan, Ann Arbor, MI, USA

**Keywords:** Influenza A virus, Baloxavir resistance mechanism, Metal modeling, MD simulation

## Abstract

Influenza A virus continues to impose a major global health and economic burden through seasonal epidemics and occasional pandemics, highlighting the critical need for continued antiviral development. As the latest addition to anti-influenza therapy, baloxavir marboxil (BXM) targets the highly conserved PA N-terminal endonuclease domain (PA_N_), blocking the cap-snatching process essential for viral transcription initiation. However, the rapid emergence of resistance mutations significantly reduces BXM susceptibility and compromises its clinical efficacy. Understanding the dynamics underlying resistance through computational modeling has been hindered by the complex electronic properties of the bimetallic catalytic center within the PA_N_ active site, posing a challenge for accurate parameterization. Therefore, in this study, we systematically benchmarked metal-parameterization strategies for molecular dynamics (MD) simulations, including non-bonded, bonded, and hybrid models, using wild-type PA_N_ in both apo and drug-bound states. Identification of reliable parameterization schemes enabled MD simulations of five clinically relevant mutants, I38T/F/M, A36V, and E23K, revealing how each reshapes the conformational landscape to modulate drug binding modes. Together, our results provide a path toward modeling complex sites in metalloenzymes and a mechanistic foundation for vulnerabilities in PA_N_ to guide structure-based optimization of next-generation inhibitors.

**TOC:** 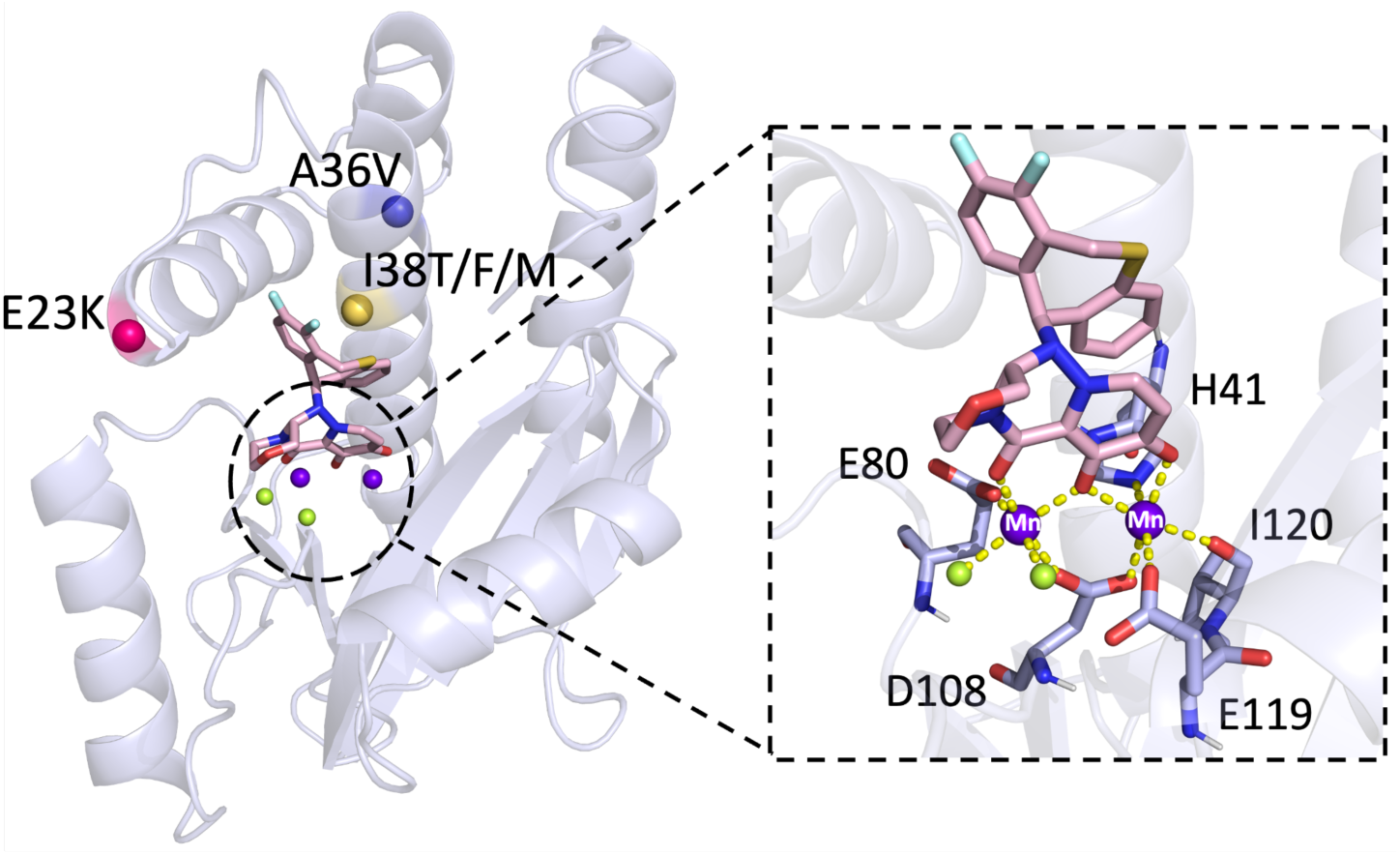

## Introduction

Influenza A virus (IAV) is a major respiratory pathogen that causes acute febrile illness in the human population and poses a persistent threat to global health through seasonal epidemics and occasional pandemics.^1, 2^ According to the World Health Organization (WHO), annual seasonal epidemics result in an estimated 3 to 5 million cases of severe illness and up to 650,000 respiratory deaths worldwide. This recurring burden is driven by the antigenic drift, which refers to the gradual accumulation of mutations in surface glycoproteins hemagglutinin (HA) and neuraminidase (NA) that alter antigenic epitopes and reduce recognition by pre-existing immunity, thereby enabling immune escape and recurrent infection.^3, 4^ Wild aquatic birds serve as the natural reservoir for IAV.^5^ However, avian-derived strains can cross the species barrier to infect humans either through antigenic shift, where the reassortment of genome segments from IAVs of diverse origins during co-infection of a permissive host, such as swine, produces viruses with novel HA and/or NA subtypes,^6^ or through gradual host adaptation.^7, 8^ Due to the lack of pre-immunity in naïve populations, these novel viruses have the potential to cause severe pandemics.^9^ Consequently, currently circulating seasonal IAVs represent the descendants of past pandemic strains that have been continuously shaped by antigenic drift.^3^ Notably, since late 2021, highly pathogenic avian influenza (HPAI) H5N1 virus of clade 2.3.4.4b has caused widespread outbreaks in the United States, affecting over 100 million poultry and expanding its host range to a wide spectrum of mammals.^10^ The virus has shown considerable evolutionary plasticity, with over 100 reassortant genotypes identified in North America.^11^ Although only 74 human infections associated with exposure to infected animals have been reported to date and sustained human-to-human transmission has not been observed, increasing mammalian adaptation and reassortment underscore the ongoing risk of cross-species transmission and potential pandemic emergence.

Presently, vaccination remains the most effective strategy against IAV, but necessitates annual updates to keep up with antigenic variation. Moreover, vaccine effectiveness varies across host populations and is limited during the early stages of an IAV pandemic.^12, 13^ Under these conditions, antivirals emerge as the most immediately effective weapon; however, their effectiveness is increasingly undermined by resistance. Early M2 blockers are now obsolete due to widespread resistance^14^ and central nervous system side effects,^15^ and neuraminidase inhibitors (NAIs) continue to encounter sporadic resistant strains despite improvements since the 2009 H1N1 pandemic.^6, 16^ This persistent challenge has catalyzed the development of baloxavir marboxil (BXM), the latest approved antiviral that introduces a novel therapeutic class as a cap-dependent endonuclease inhibitor (CENI). BXM is intracellularly hydrolyzed to its active form, baloxavir acid (BXA), which targets the N-terminal endonuclease domain of PA (PA_N_) within the RNA-dependent RNA polymerase (RdRp), a heterotrimer composed of PA, PB1, and PB2, that is responsible for transcription and replication of the eight-segment, negative-sense IAV RNA genome.^17, 18^ RdRp initiates transcription via a unique “cap-snatching” mechanism in which PB2 binds the 5′ m^7^G cap of host pre-mRNAs and the metal-ion-dependent PA_N_ cleaves them approximately 10-13 nucleotides downstream to generate capped primers for PB1-mediated mRNA synthesis.^9, 19^ By targeting PA_N_, BXA effectively blocks primer generation, thereby halting viral replication at its source. Despite its clinical efficacy, with a single dose achieving symptom relief comparable to NAI oseltamivir while demonstrating superior viral load reduction and shortening the median duration of infectious virus detection to 24h, BXM has not escaped the emergence of resistance.^20, 21^ Multiple clinical studies^22, 23^ and global surveillance efforts^24, 25^ have identified amino acid substitutions at position 38 of the PA subunit (I38T/F/L/M/N/S/V) as the key determinant of resistance to BXM, with I38T being the most severe, resulting in an approximately 11- to 124-fold increase in EC_50_ for viral replication.^26^ Additional substitutions, such as E23K/G, A36V, A37T, and E199G, have also been reported, but occur much less frequently and confer only modest reductions in BXA susceptibility. Current resistance mutations, while facilitating viral escape, concurrently compromise in vitro replication fitness and limit widespread transmission. However, serial passaging has revealed that I38L variants can acquire potential compensatory substitution, D394N, partially offsetting the associated fitness cost.^27^ This pattern mirrors the evolutionary history of NAI oseltamivir resistance, in which compensatory substitutions can restore replicative fitness and enable widespread global prevalence.^28^ Collectively, these parallels suggest that BXM-resistant viruses may similarly acquire compensatory mutations and achieve broader circulation, highlighting the importance of elucidating resistance mechanisms to guide the rational design of next-generation antivirals with improved resilience to resistance.

Current structural studies of BXM resistance have predominantly focused on the I38T substitution, where reduced BXA efficacy is attributed to weakened van der Waals (VDW) packing and an induced-fit rotameric rearrangement of Thr38 upon BXA binding.^23, 29^ However, static crystal structures are unable to capture the full spectrum of resistance mechanisms, as protein function is intrinsically governed by the dynamic distribution of conformational ensembles. This is exemplified by the distal mutation A36V, whose crystal structure shows no significant deviation from the wild-type (WT), suggesting a potential resistance mechanism driven by altered protein dynamics rather than direct disruption of inhibitor binding.^28^ Limited molecular dynamics (MD) simulations^30–34^ have been conducted on PA_N_, but these studies relied on automatically generated metal parameters without thorough evaluation and primarily focused on Ile38 substitution, leaving the resistance mechanisms underlying other key clinically relevant variants largely unresolved. Given that BXA exerts its inhibitory effect by chelating two Mn^2+^ ions in the active site, critical consideration of the bimetallic center is necessary for reliable evaluation of how resistance mutations influence PA_N_ dynamics and BXA binding.

Computational modeling of metal centers in metalloproteins is particularly challenging. The high charge density of metal ions induces strong polarization and charge transfer, rendering the electronic environment of coordinating residues highly sensitive to local conformational fluctuations. For transition metals, such as the divalent Mn ions in PA_N_, partially filled d orbitals further impose strong directional preferences on coordination geometry.^35, 36^ Although polarizable force fields and hybrid quantum mechanics/molecular mechanics (QM/MM) methods can describe metal centers with high accuracy, their increased computational costs and methodological complexity prohibit routine long-timescale sampling of complex ligand binding modes and global conformational dynamics.^37^ Consequently, non-polarizable models remain the dominant strategy for metalloprotein parameterization.^38^ The simplest nonbonded model represents metal ions as point charges interacting through Coulombic and 12-6 Lennard-Jones terms, allowing coordination number changes and ligating residue exchange, but often struggles to reproduce accurate coordination geometries due to its neglect of polarization, charge transfer, and coordination directionality.^39^ The 12-6-4 Lennard-Jones potential further refines the nonbonded framework by adding a C_4_ term to approximate ion-induced dipole interactions.^40^ This term adds a pairwise coefficient scaled by r^-4^. Conversely, bonded models preserve local geometry through explicit covalent terms at the cost of artificially restricting ligand exchange and coordination variability. To bridge this gap, hybrid approaches strategically combine bonded restraints for coordinating amino acids with nonbonded treatments for labile ligands, effectively balancing stability with sampling flexibility.^37, 41^ Due to this complexity, it is not always clear which model is appropriate for a given system, especially in the presence of small-molecule ligands, where little information regarding metal-ligand interactions is available.

Therefore, to represent PA_N_ dynamics with high accuracy, we thoroughly investigated each of these non-polarizable metal modeling strategies in both the WT apo and BXA-bound systems. Tracking the coordination geometry over 500 ns of MD simulations revealed that both the 12-6 and 12-6-4 nonbonded models successfully maintained the appropriate architecture for the apo bimetallic site. However, the bonded model was necessary to model the BXA-bound site, given the lack of accurate ligand-specific polarization treatment in current nonbonded models, highlighting an important avenue for future development. Nevertheless, these strategies enabled modeling of five clinically relevant PA_N_ variants, including I38T/F/M, A36V, and E23K. By performing three independent replicates of 500 ns MD simulations for each system, our study uncovered that the I38 substitutions (T/M/F) reduce BXA efficacy by disrupting the stable Ile38-mediated hydrophobic anchor, which directly destabilizes the bound BXA. In contrast, A36V acts primarily in the apo state, where local α-helical unwinding promotes sampling of more open pocket conformations and perturbs pre-binding organization. Furthermore, the E23K mutation rewires the PA_N_ α2-helix electrostatic network, weakening key α2-associated interactions that stabilize BXA. Together, these findings provide both a practical framework for modeling complex metalloenzymes and a dynamic structural basis for understanding BXA resistance, further guiding the design of next-generation CENIs.

## Methods

### Homology modeling

All systems, including apo and BXA-bound WT PA_N_ as well as the I38T/M/F, E23K, and A36V mutants from the A/California/04/2009 (H1N1) strain, were constructed based on available crystal structures obtained from the Protein Data Bank (PDB)^42^ (**Table 1**). Variants lacking experimental structures were generated by introducing point mutations into the corresponding WT templates using tleap from AmberTools24.^43^ To reconstruct the missing flexible loop (residues 51-72) truncated during crystallization, the FASTA sequence of PA_N_ was retrieved from UniProt (accession C3W5S0) and used to identify homologous templates (PDB IDs: 4LN7, 4M4Q, and 9DOJ) via BLAST searches^44^ against the PDB. Using MODELLER 10.6,^45^ 100 candidate models were generated per system. During this process, reconstruction was restricted to the loop region, while the remainder of the protein was kept fixed. Models were evaluated using the Discrete Optimized Protein Energy (DOPE) score,^46^ a statistical potential for model quality, and the lowest-scoring structure was selected as the final model.

**Table 1.** Structural information for apo and BXA-bound PA_N_ systems.

| PA<sub>N</sub> Systems | Apo | BXA-bound |
| --- | --- | --- |
| WT | 8T5Z | 7K0W |
| I38T | 8T67 | 6FS7 |
| A36V | Modeled from 8T5Z | 8T5V |
| E23K | 8VGZ | Modeled from 7K0W |
| I38F | Modeled from 8T5Z | Modeled from 7K0W |
| I38M | Modeled from 8T5Z | Modeled from 7K0W |

### Bimetallic active site parameterization

#### Non-bonded model

Two nonbonded models, 12-6 and 12-6-4, were evaluated for modeling the two divalent Mn ions in the PA_N_ active site. The 12-6 nonbonded model treats the interactions between metal and other particles entirely through non-covalent terms, with van der Waals interactions described by the 12-6 Lennard-Jones (LJ) potential and electrostatic interactions described by the classical Coulomb potential. The 12-6 normal usage parameter set^39^ optimized for the Amber TIP3P water model was utilized for the two Mn^2+^ ions.

The 12-6-4 nonbonded model extends the traditional 12-6 LJ potential by introducing an additional C_4_ (pairwise coefficient scaled by r^-4^) term to account for ion-induced dipole interactions. The 12-6-4 parameters between Mn^2+^ and the TIP3P water model are from Li and Merz.^40^ For coordinating amino acids, C_4_ terms between Mn^2+^ and imidazole nitrogen and between Mn^2+^ and carboxylate oxygen were taken from Li et al.^47^ and Jafari et al.,^48^ respectively, both of which were developed specifically for TIP3P. The unpublished C_4_ terms between Mn^2+^ and hydroxide oxygen, and between hydroxide oxygen and the TIP3P water were provided by the Merz group. Specifically, the C_4_ term between Mn^2+^ and OH^-^ was parameterized based on the logK of 3.4^49^ using umbrella sampling, following the approach described in refs 47 and 48. The C_4_ term between OH^-^ and TIP3P water was parameterized based on the hydration free energy of OH^-^ as −105.0 kcal/mol^50^ using thermodynamic integration, following the approach described in ref 40. For coordinating atom types without established C_4_ parameters, the carbonyl oxygens of Ile120 backbone and BXA were approximated using the carboxylate oxygen parameter, whereas the deprotonated BXA enolate oxygen was approximated using the hydroxide oxygen parameter. All the C_4_ terms were added via ParmEd. Care was taken to ensure compatibility with the GPU-accelerated PMEMD (CUDA) engine by verifying that all C_4_ donors possessed non-zero C_4_ values and their atom types included explicit charge designations (i.e., “+” or “-”).

In both nonbonded models, coordinating waters were described using the TIP3P model. The bridging hydroxide anion in apo systems and the BXA in bound systems were parameterized separately, with RESP charges^51^ derived from Hartree-Fock/6-31G(d) calculations in Gaussian16^52^ and force field parameters assigned from the GAFF2 force field.^53^

#### Bonded model

The bonded model for the binuclear manganese center was constructed using the MCPB.py^54^ program. Initially, two models were generated for QM calculations at the B3LYP/6-31G(d) level to derive the force field parameters. The sidechain model consisted of the two Mn^2+^ ions, coordinating water molecules, bridging hydroxide or BXA, and the truncated coordinating amino acid fragments. For amino acids coordinating through side-chain atoms, the side chains were retained and capped with a methyl group, whereas amino acids coordinating through the backbone were represented by an ACE-NME fragment. The large model included the two Mn^2+^ ions, coordinating water, bridging hydroxide or BXA, with the full coordinating residues capped with acetyl and N-methylamide groups. Considering the coordination environment of the binuclear manganese center in PA_N_ and its known preference in biological systems,^55, 56^ the two Mn^2+^ ions were modeled in a high-spin antiferromagnetic coupling (AFM) state. Specifically, the geometry was first optimized at the high-spin ferromagnetic (FM) state to obtain a reasonable initial geometry. Subsequently, a broken-symmetry spin configuration guess was generated using the fragment-based approach implemented in Gaussian to describe the AFM singlet state. In the apo system, each Mn^2+^ ion together with its directly coordinating residues was defined as an individual fragment, whereas the bridging hydroxide and Asp108 were treated as separate fragments. The same fragmentation strategy was applied to the BXA-bound system, with BXA and the bridging Asp108 treated as independent fragments. After full geometry optimization, frequency calculations were performed on the sidechain model to obtain bond and angle force constants using the Seminario method, while the electrostatic potential was computed on the large model to derive partial charges via RESP fitting using the ChgModB scheme. Notably, since two water molecules coordinate the same Mn^2+^ ion in our systems, a standard bonded representation would introduce artificial 1-4 interactions, potentially causing structural distortions.^57^ Furthermore, to preserve the dynamic flexibility of the ligands, interactions were maintained via harmonic restraints rather than fixed bonded interactions. This approach maintains coordination distances near their equilibrium, while allowing greater flexibility in the coordination geometry.

#### Hybrid model

To preserve the coordination geometry while allowing ligand flexibility and exchange, a hybrid modeling scheme combining features of both bonded and non-bonded models was employed for the binuclear manganese center. In contrast to the bonded model, RESP charge fitting based on the calculated electrostatic potential of the large model was applied only to the Mn^2+^ ions and the coordinating amino acids, whereas the partial charges of the coordinating waters and BXA were fixed to values derived from the non-bonded model. In addition, the harmonic restraints between the Mn^2+^ ions and the coordinating atoms of the water molecules and BXA, which would otherwise be introduced in a bonded representation, were removed. All other parameters and settings were kept consistent with those of the bonded model.

#### MD simulation

System preparation was facilitated by the tLEaP program in AmberTools24. The Amber ff14SB force field^58^ was applied to all proteins, while specialized force field parameters for the bimetallic active center were assigned according to the corresponding metal modeling strategies described above. His41 was modeled in the HID state to accommodate metal coordination. Each system was solvated in an isometric TIP3P water box, ensuring a minimum distance of 13 Å between the solute and the box edges. The systems were neutralized by adding Na⁺ ions, followed by the addition of 0.15 M NaCl to mimic physiological ionic strength. Energy minimization was then conducted for 50,000 cycles with restraints applied to the solute, gradually reduced from 50 to 0 kcal·mol^-1^·Å^-2^. The systems were then heated from 0 to 310 K over 350 ps under the NVT ensemble with restraints decreased from 20 to 0 kcal·mol^-1^·Å^-2^, followed by a 1 ns equilibration under the NPT ensemble and a 500 ns production run. Throughout all simulations, periodic boundary conditions were employed. Long-range electrostatic interactions were treated using the particle mesh Ewald method^59^ with a non-bonded cut-off of 10 Å. The temperature was maintained at 310 K using a Langevin thermostat with a collision frequency of 1.0 ps^-1^. The SHAKE algorithm was applied to constrain bonds involving hydrogen atoms. Three independent replicas were performed for each system to ensure statistical reliability.

#### Trajectory analysis

All trajectory analyses were performed using the cpptraj module of AmberTools24. To characterize the coordination geometry of the bimetallic active center, the radial distribution function (RDF) was computed from the trajectories with a resolution of 0.01 Å, with densities normalized to the average simulation volume. The first minimum of each RDF was used to define the boundary of the first coordination shell and as the integration limit for calculating the average coordination number (CN). The same cutoff was then used to calculate contact frequencies of coordinating atoms with the two Mn^2+^ ions. Principal component analysis (PCA) was performed using backbone atoms to characterize the dominant collective motions of the protein. To separate loop-driven motions from intrinsic core conformational changes, PCA was carried out separately for the whole protein and for the structured core excluding the flexible 51-72 loop. For each analysis, all systems were used to construct and diagonalize a mass-weighted coordinate covariance matrix, and the trajectories were projected onto the resulting common principal component space to compare conformational sampling across systems. Kernel density estimation (KDE) was subsequently applied to the projections along the leading principal components to visualize the conformational probability density using Gaussian kernels, eight contour levels, and a density threshold of 0.05. Density-based clustering was also performed in the reduced PCA space using the mean-shift algorithm implemented in scikit-learn, with bandwidth automatically estimated from the PCA projections. For each cluster, the frame closest to the cluster center was selected as the representative structure. Pocket volumes were calculated using POVME 3^60^ with a 1.0 Å grid spacing. All structures were first aligned to the representative WT BXA-bound structure obtained from loop-excluded PCA clustering, and the point inclusion sphere was defined from the same active-site pocket center with a 9.0 Å radius. Protein-ligand interactions were characterized across the trajectories using ProLIF.^61^ ProLIF uses RDKit molecular representations and built-in SMARTS-based atom selections to identify relevant chemical features, and assigns interactions based on predefined distance and angle criteria. The resulting interaction fingerprints were used to quantify the occupancy of residue-level protein-ligand contacts across the trajectories.

## Results and Discussion

### Both 12-6 and 12-6-4 nonbonded models preserve the apo PA_N_ bimetallic coordination geometry

Several crystal structures have resolved the intrinsic coordination geometry of the PA_N_ binuclear manganese center by removing the flexible loop spanning residues 51-72 that previously introduced crystal packing artifacts.^29, 62, 63^ Both Mn^2+^ ions adopt well-defined octahedral coordination geometries. In the apo state, the two Mn^2+^ ions are bridged by a hydroxide anion, with the first manganese (M1) further coordinated by His41, Asp108, Glu119, Ile120, and one water molecule, while the second manganese (M2) is coordinated by Glu80, Asp108, and three water molecules (**Figure 1A**). In the BXA bound state, three oxygen atoms from BXA form a tridentate chelate with the two Mn^2+^ ions, replacing the bridging hydroxide anion, and two coordinating water molecules, preserving the overall octahedral framework **(Figure 1B)**. To establish an accurate description of this catalytic bimetallic center in a physiological context, we first reconstructed the missing loop via homology modeling and benchmarked different metal parameterization strategies for conducting MD simulations of full-length WT PA_N_ in both apo and BXA-bound states. The ultimate goal of this study is to elucidate how resistance mutations perturb drug binding, therefore a nonbonded representation of the metal center was prioritized, as it avoids imposing geometric constraints, allowing BXA and coordinating residues to adjust dynamically during simulations.

**Figure 1.**
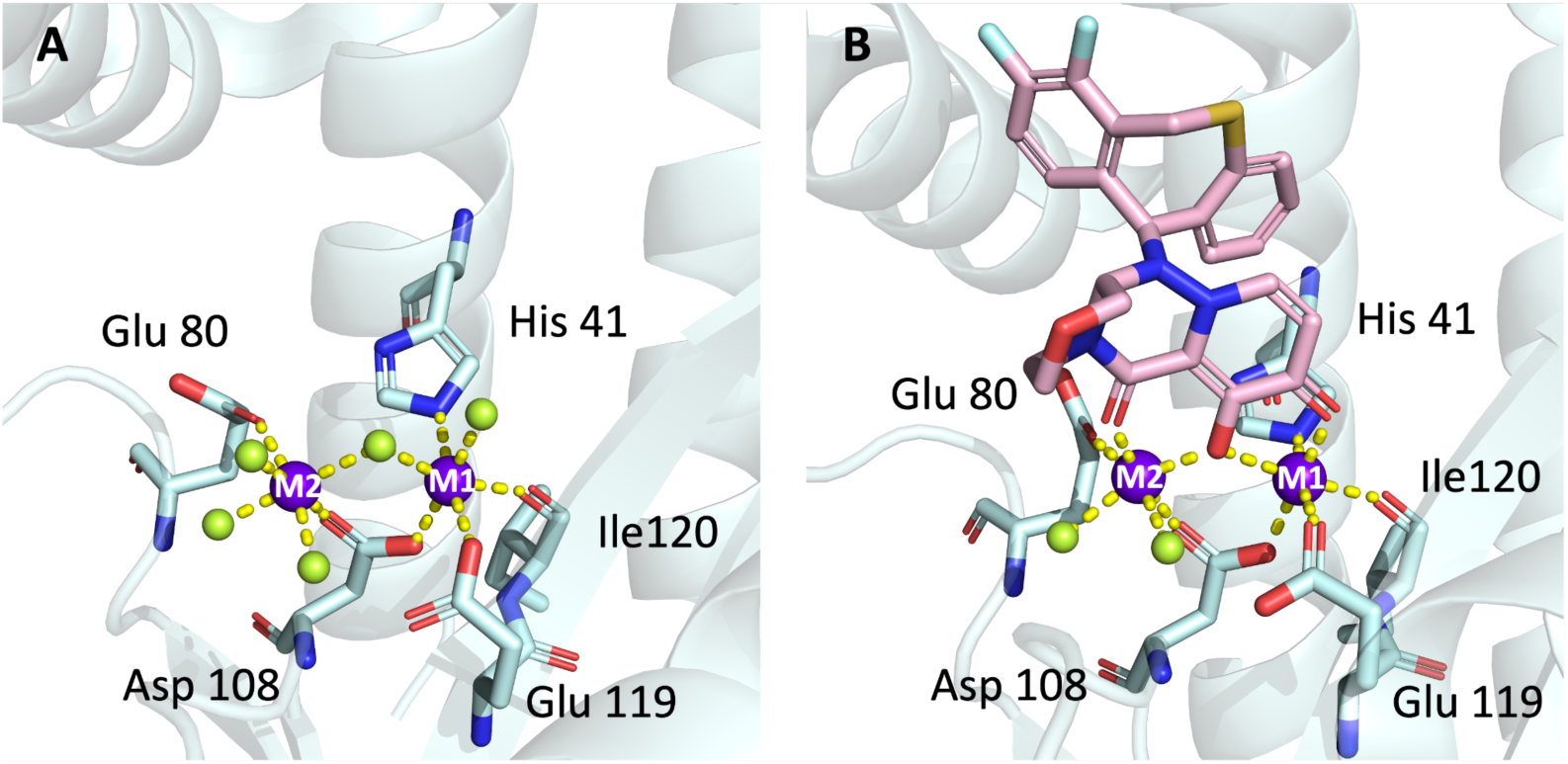
Coordination geometry of the PA_N_ bimetallic active center in apo and BXA-bound states. **(A)** Apo PA_N_ active site (PDB ID: 8T5Z). **(B)** BXA-bound PA_N_ active site (PDB ID: 7K0W).

For the apo system, we first evaluated the 12-6 nonbonded model by monitoring the overall system stability and metal coordination geometry throughout 500 ns of simulation. Although motions of the flexible loop occasionally increased the overall backbone RMSD, the structured core remained highly stable at around 1.0 Å, while both Mn^2+^ ions exhibited minimal fluctuations with RMSD largely below 0.5 Å throughout the trajectories (**Figure 2A**). To characterize the metal coordination environment, we computed the radial distribution function (RDF) for each Mn^2+^ ion and potential coordinating atoms. Integration of the first RDF peaks yielded average coordination numbers, with the protein, water, and total first-shell contributions all aligning with theoretical expectations (**Figure 2B**). By utilizing the first-shell RDF minima as distance cutoffs, we confirmed that both Mn^2+^ ions remained stably coordinated by their native residues, with contact frequencies approaching 100%. Furthermore, frame-by-frame contact analysis revealed that M1 and M2 maintained the ideal hexacoordinated geometry for 99.61% and 100% of the simulation time, respectively. Next, we removed crystallographic waters and tested whether waters from the solvent could spontaneously coordinate. In this setup, solvent molecules rapidly reorganized within the first few nanoseconds to restore the coordination environment, after which both the coordination geometry and protein structure remained stable. Collectively, these results indicate that electrostatic and van der Waals interactions alone are sufficient to stabilize the binuclear manganese center in the apo system and the spontaneous coordination recovery corroborates the physical realism of the 12-6 non-bonded parameters (**Figure 2C**).

**Figure 2.**
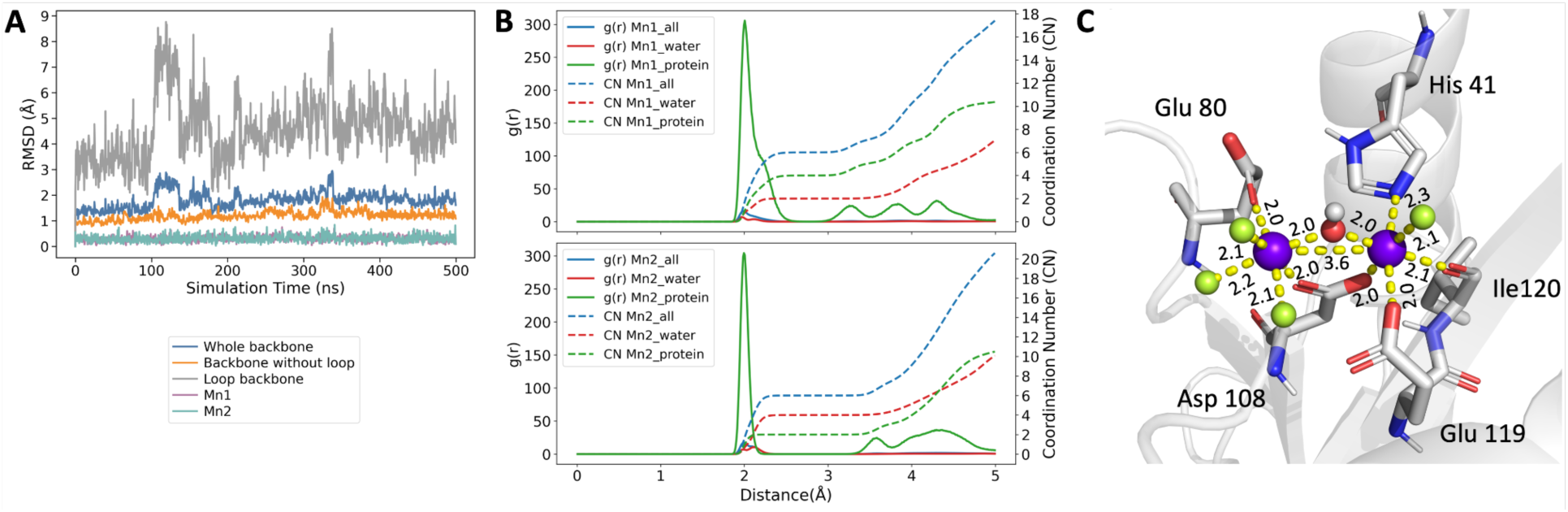
Stability and bimetallic coordination geometry of apo PA_N_ during 500 ns MD simulation using the 12-6 nonbonded model. **(A)** RMSD calculations for the full protein backbone, protein backbone excluding the flexible loop, flexible-loop backbone only, and the two Mn^2+^ ions. All trajectories were first aligned using the protein region excluding the flexible loop for RMSD calculation. **(B)** RDF plots for each Mn^2+^ ion were calculated using either three groups of potential coordinating atoms, including all system atoms, water and hydroxide oxygen atoms, and protein atoms. The sharp first peak corresponds to the first coordination shell, whereas the broader peaks at longer distances represent outer-shell or non-coordinating neighboring atoms. **(C)** Representative structure showing the coordination geometry of the bimetallic center. Average coordination distances are labeled in Å. The bridging hydroxide ion positioned between the two Mn^2+^ ions is shown with oxygen in red and hydrogen in white, Mn^2+^ ions shown in purple, and coordinating water molecules shown in limon.

We next tested the 12-6-4 nonbonded model in the apo PA_N_ system, which incorporates ion-induced dipole interactions through an additional C_4_/r^-4^ term. With this model, the overall PA_N_ structure remained stable, and both Mn^2+^ ions retained the ideal hexacoordinated geometry for around 99.9% of the simulation time **(Figure S1)**. Interestingly, unlike in the 12-6 simulation, one of the three M2-coordinating waters (W4) underwent continuous exchange with solvent water molecules throughout the 500 ns trajectories. This behavioral difference likely arises because the standard 12-6 model tends to bind coordinating waters within a deeper potential well than the 12-6-4 model. In contrast, the C_4_/r^-4^ term in the 12-6-4 model modulates the metal-water interaction, increasing the inherent solvent kinetic flexibility of the metal coordination shell.

### Nonbonded models fail to accurately represent the BXA-bound bimetallic center

For the BXA-bound system, the coordination geometry began to distort within the first few nanoseconds when using the 12-6 nonbonded model. BXA gradually shifted away from the Mn^2+^ ions, resulting in loss of key metal-ligand interactions and replacement of coordinating atoms by solvent molecules or nearby residues, while each Mn^2+^ ion still largely retained a six-coordinate geometry **(Figure 3A)**. This behavior highlights the limited transferability of the water-derived 12-6 parameters when applied to a complex ligand-bound metal center. The instability likely arises from neglecting the polarization effects critical for divalent metal ions, resulting in a failure to accurately capture the short-range electrostatic balance required for strong metal-ligand interactions and ultimately destabilizing the coordination geometry.

**Figure 3.**
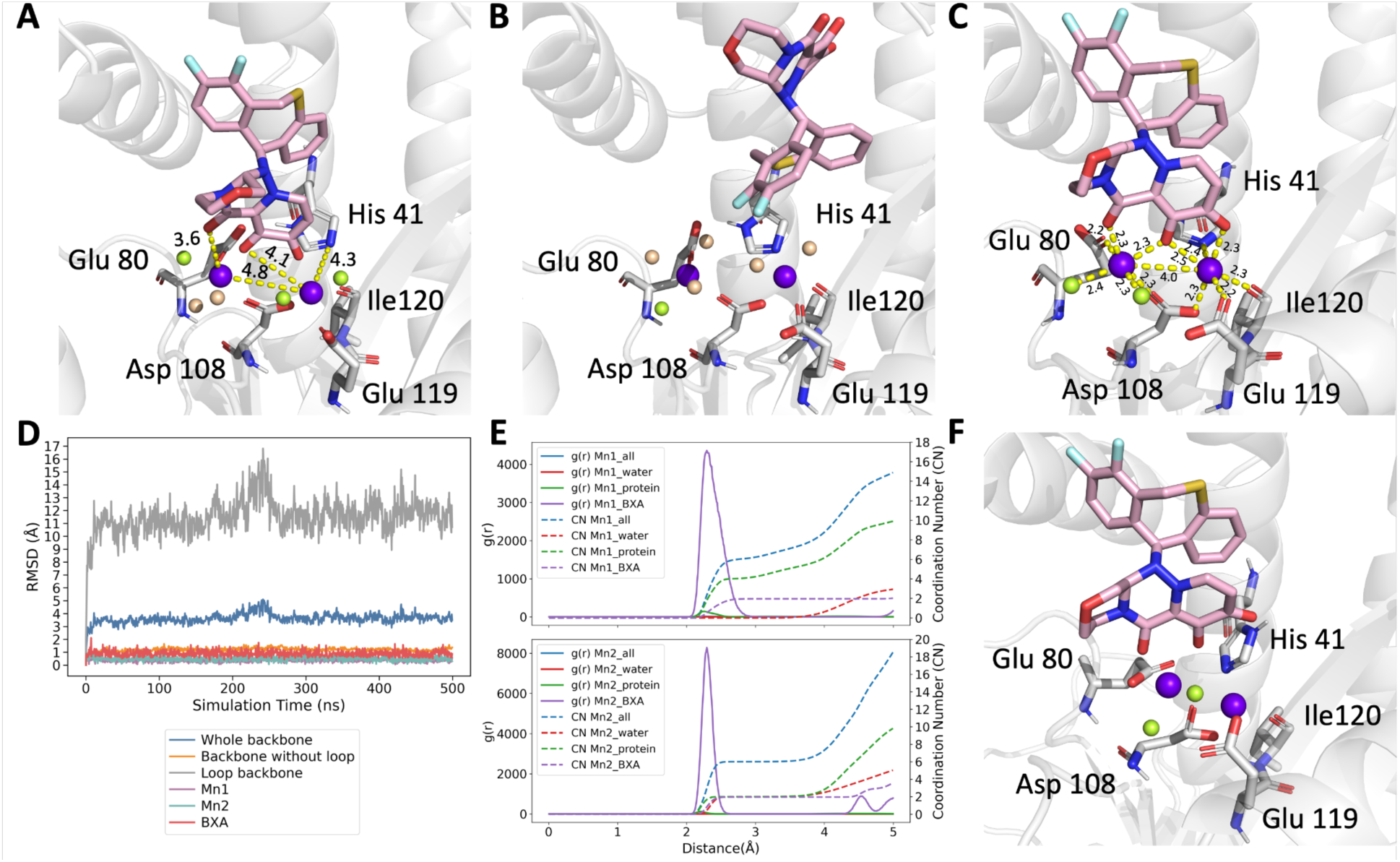
Performance of different metal modeling strategies in the BXA-bound PA_N_ active site. **(A-B)** Snapshots from BXA-bound systems parameterized with the 12-6 nonbonded model at 100 ns and the 12-6-4 nonbonded model at 30 ns, respectively. Both strategies failed to maintain the native BXA-bound coordination geometry. Native coordinating waters are shown in limon, and newly coordinated waters are shown in wheat. **(C)** Representative bonded-model structure showing preserved coordination geometry, with average coordination distances labeled in Å. **(D)** RMSD profiles of the BXA-bound system parameterized with the bonded model, including the protein backbone, backbone excluding the flexible loop, loop backbone, BXA, and the two Mn^2+^ ions. **(E)** Average RDFs and coordination numbers for the bonded model, showing first-shell coordination consistent with the expected BXA-bound metal environment. **(F)** Snapshot from the BXA-bound system modeled using the hybrid model at 6 ns, showing failure to maintain the native coordination geometry. Native coordinating waters are shown in limon.

Therefore, we further tested the 12-6-4 nonbonded model, which introduces ion-induced dipole interactions by manually assigning C_4_ parameters for the interactions between the Mn^2+^ ions and the relevant coordinating atom types, as detailed in the Methods section. This model also failed to sustain a stable coordination environment. Within approximately 30 ns of the simulation, the BXA completely dissociated from the metal center and diffused out of the active site **(Figure 3B)**. This unphysical behavior likely stems from the unavailability of C_4_ parameters specifically calibrated for the interactions between Mn^2+^ and the carboxylate or deprotonated enolate oxygens of the BXA. The reliance on approximate C_4_ values proved insufficiently precise to properly anchor the ligand within the bimetallic site.

Finally, we employed the bonded-model strategy to stabilize the binuclear manganese center in the BXA-bound system. Force constants and RESP charges were determined using quantum mechanics (QM) calculations (see Methods for details). Because one of the Mn^2+^ ions is coordinated by multiple water molecules simultaneously, directly applying a standard bonded representation would introduce artificial 1-4 interactions between these waters. Since standard water models lack hydrogen VDW parameters, the 1-4 VDW interactions are effectively zero, whereas the attractive 1-4 electrostatic interactions remain, leading to unphysical distortion of the local geometry. Therefore, the bonded model was implemented through harmonic distance restraints rather than fixed bonded interactions. By applying the QM-derived bond force constants to maintain the metal-ligating atom equilibrium distances, this strategy should stabilize the complex without over-restricting it, thereby preserving a necessary degree of structural flexibility. The simulations showed that both the protein backbone and the bimetallic center remained stable **(Figure 3C-E)**, with M1 and M2 retaining their expected coordination geometries for 92.06% and 99.95% of the simulation time, respectively.

We additionally tested a hybrid modeling strategy, in which harmonic restraints were applied only between the Mn^2+^ ions and coordinating amino acid residues, while coordinating waters and BXA were treated as fully nonbonded to allow solvent exchange and unrestricted BXA motion. However, this approach was insufficient to stabilize the BXA-bound bimetallic center **(Figure 3F)**. This outcome was expected because the RESP-derived partial charges assigned to the Mn^2+^ ions in the bonded model are lower than their formal +2 oxidation state. Consequently, interactions between the metal ions and the atoms treated as nonbonded become underestimated, ultimately destabilizing the coordination network.^64^

To maintain a consistent metal-modeling strategy across both the apo and BXA-bound systems, we attempted to parameterize a bonded model for the apo state. However, the apo bimetallic center failed to preserve its experimentally observed coordination geometry during QM optimization. We hypothesize that, absent the geometric constraints imposed by the tridentate BXA inhibitor, the apo coordination environment is intrinsically more flexible and thus challenging to stabilize using the employed basis set. To overcome this issue, heavy atoms were frozen during the subsequent optimization, which ultimately enabled the generation of reasonable bonded-model parameters for the apo system and preserved the native coordination geometry **(Figure S2)**. We also evaluated the hybrid strategy for the apo system in which harmonic restraints were applied only between the Mn^2+^ ions and coordinating amino acids, while coordinating waters remained fully exchangeable. Under this condition, the metal center also remained stable, with coordinating water molecules undergoing continuous exchange with solvent molecules while preserving the overall coordination geometry **(Figure S3)**. Despite these promising results, deriving force field parameters by artificially freezing heavy atoms is not preferred. Given that the straightforward 12-6 nonbonded model already performs exceptionally well for the apo state, we ultimately employed the 12-6 nonbonded model for all subsequent apo simulations. Future studies may further improve apo bonded models by employing higher levels of QM theory to optimize the apo metal center without artificial constraints or tuning VDW parameters to stabilize hybrid representations of the bound system; however, such developments fall beyond the scope of the present work. Collectively, benchmarking of various metal modeling strategies across the apo and BXA-bound WT PA_N_ systems demonstrated that nonbonded models were sufficient to stabilize the apo metal center, but failed to accurately maintain the more complex BXA-chelated coordination network in the bound state due to a lack of accurate system-specific polarization treatments. Because the bonded model was the only strategy capable of reliably preserving the bimetallic center, it was adopted for all subsequent BXA-bound simulations.

### WT PA_N_ establishes the baseline conformational dynamics and BXA binding mode

Having established optimal modeling strategies to treat the binuclear manganese center, we conducted MD simulations to elucidate the molecular mechanisms underlying BXA resistance driven by five PA_N_ mutants. For both the WT and mutant systems, three independent 500 ns replicas were simulated in both the apo and BXA-bound states.

Structurally, the native PA_N_ consists of seven α-helices, a mixed five-stranded β-sheet, and a flexible loop spanning residues 51-72. Throughout the simulations, although motions of the reconstructed loop contributed to elevated overall protein backbone RMSDs and occasional large fluctuations, the structured core region remained highly stable in both apo and BXA-bound systems, with backbone RMSDs centered around 1.0 Å after excluding the flexible loop (**Figure 5A**). Consistently, RMSF analysis identified the 51-72 loop as the dominant source of conformational flexibility in both WT states (**Figure 5B**). The active site architecture and binding network were also well-maintained. In agreement with the benchmarked metal-modeling strategy described above, both Mn^2+^ ions remained stably coordinated and preserved the expected octahedral coordination geometries throughout the trajectory replicas. BXA remained tightly bound within the catalytic pocket throughout the simulation in the drug-bound system (**Figure 5A**), adopting a butterfly-like conformation in which the oxazino-pyridotriazin-dione moiety serves as the anchor domain coordinating the bimetallic center, while the difluoro-dihydro-dibenzothiepine wing functions as the specificity domain, extending into an adjacent hydrophobic subpocket to confer binding selectivity and stabilize a defined binding pose through extensive hydrophobic and aromatic interactions **(Figure 5C)**. Analysis of protein-BXA interaction frequency across the trajectories revealed that Ile38 acted as the dominant hydrophobic anchor for the specificity domain, maintaining persistent hydrophobic contacts throughout the simulations. His41 and Tyr24 also engaged this wing through nearly continuous hydrophobic contacts, while Lys34 and Met21 maintained frequent contact, with occupancies of 82.5% and 767.6%, respectively. Among aromatic interactions, Tyr24 acted as the primary π-stacking residue, exhibiting π-π interactions with the specificity domain in 56.5% of the trajectory. In contrast, His41 and Phe105 contributed more transient π interactions, with His41 interacting with the specificity domain for 23.3% of the trajectory and Phe105 interacting with the anchor domain for 13.9% (**Figure 5D**).

**Figure 5.**
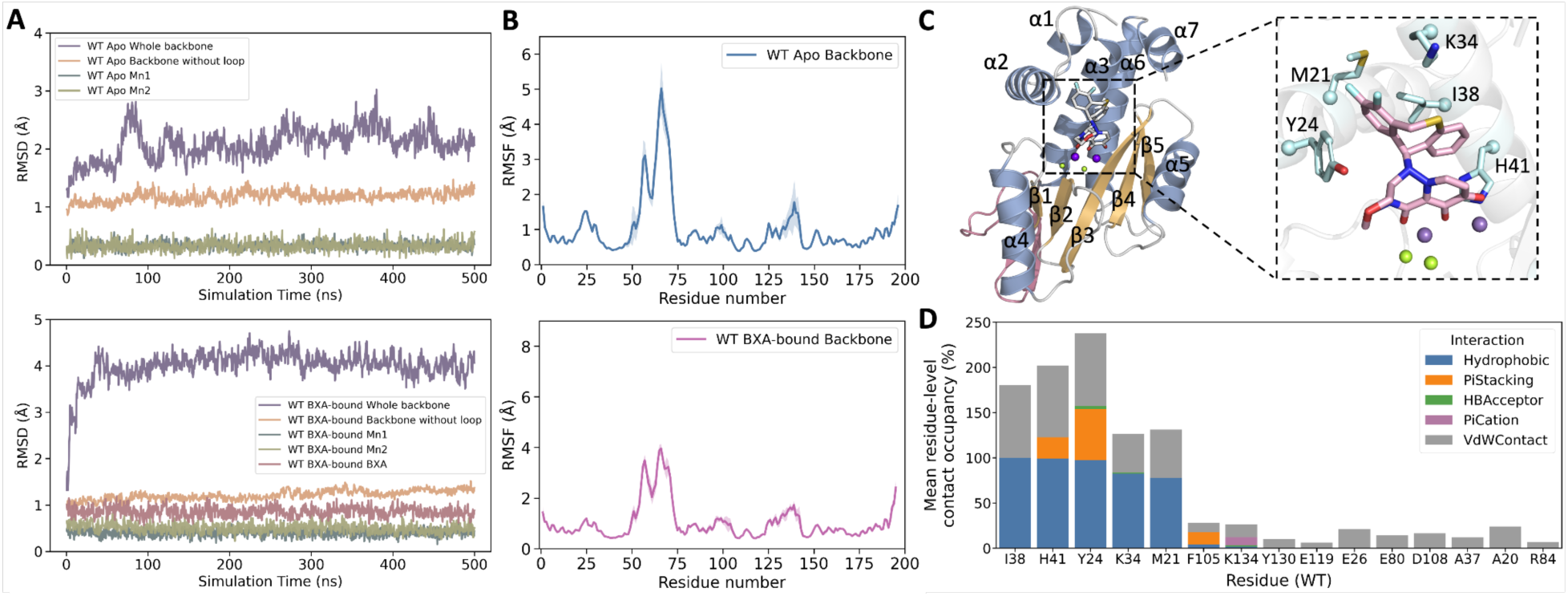
Summary of WT PA_N_ dynamics and BXA binding interactions in both apo and BXA-bound simulations. **(A)** RMSD profiles of the WT apo and BXA-bound systems. For each state, RMSDs were calculated for the full protein backbone, protein backbone excluding the flexible loop, two Mn^2+^ ions, and BXA where applicable. **(B)** Residue-level RMSF profiles of WT apo and BXA-bound PA_N_. Shaded regions indicate standard deviations across three replicas. **(C)** Representative BXA-bound WT PA_N_ structure, with α-helices shown in blue, β-strands in wheat, and the flexible loop in pink. Mn^2+^ ions are shown in purple, and coordinating waters are shown as lime spheres. The inset highlights the BXA binding pose and key interacting residues. **(D)** Residue-level BXA interaction occupancy across the trajectories. Each colored segment represents the occupancy of a specific interaction type for a given residue, defined by the presence of at least one corresponding atom-level contact in a frame. Thus, 100% occupancy indicates that the residue-level interaction was present in all analyzed frames. Because interaction types were quantified independently and may occur simultaneously, the cumulative stacked occupancy for a residue can exceed 100%.

Overall, these results demonstrate the robustness of the simulation protocols and establish a comprehensive WT reference baseline for subsequent comparisons with resistance-associated mutants. In the following sections, we focus on the primary mutant-specific changes that provide mechanistic insight into BXA resistance.

### Loss of native hydrophobic networks and destabilization of BXA binding drive resistance in I38 mutants

Ile38 is a major hydrophobic anchor for BXA, and substitutions at this position represent the most frequent resistance-associated changes. Among them, I38T is recognized as the dominant BXA resistance marker due to its pronounced impact on BXA efficacy.^65^ In the apo and BXA-bound states, the overall protein backbone RMSD, RMSF, and PCA differences between mutants and WT were mostly negligible (**Figure S4-5**). The largest impact of the Ile38 mutations was observed at the binding site, with increased BXA RMSD and RMSF, distinct interaction profiles, and binding modes (**Figure 6A-D**). For I38T, the RMSF of BXA revealed elevated fluctuations primarily at the specificity domain (**Figure 6B**). Contact frequency analysis attributed this increased mobility to the disruption of the native I38-centered hydrophobic anchor network. Specifically, replacing the hydrophobic isoleucine with a smaller, polar threonine completely abolished the hydrophobic interaction. This loss was accompanied by a decrease in the neighboring Lys34 hydrophobic contact from 82.5% to 72.3% (**Figure 6C**). Structurally, the BXA binding pose differs from WT, with the difluorinated aromatic ring shifting deeper into the pocket toward T38 (**Figure 6D**). This repositioning brought the inhibitor closer to Met21, consistent with the compensatory increase in Met21 hydrophobic occupancy from 77.6% in WT to 94.8% in I38T. In addition, subtle upward displacement of the α5 helix, together with closer positioning of the heteroaromatic ring of BXA anchor domain, provides a plausible structural basis for the increased low-frequency Lys134 interactions, while π-stacking between this same ring and Phe105 was nearly lost, decreasing from 13.9% in WT to almost undetectable levels in I38T (**Figure 6C-D**). Collectively, consistent with previous studies,^23, 29, 30^ our results indicate that the reduced BXA efficacy of I38T primarily arises from disruption of the native hydrophobic interaction between Ile38 and the difluoro-dihydro-dibenzothiepine wing of BXA, thereby weakening stabilization of the inhibitor within the binding pocket.

**Figure 6.**
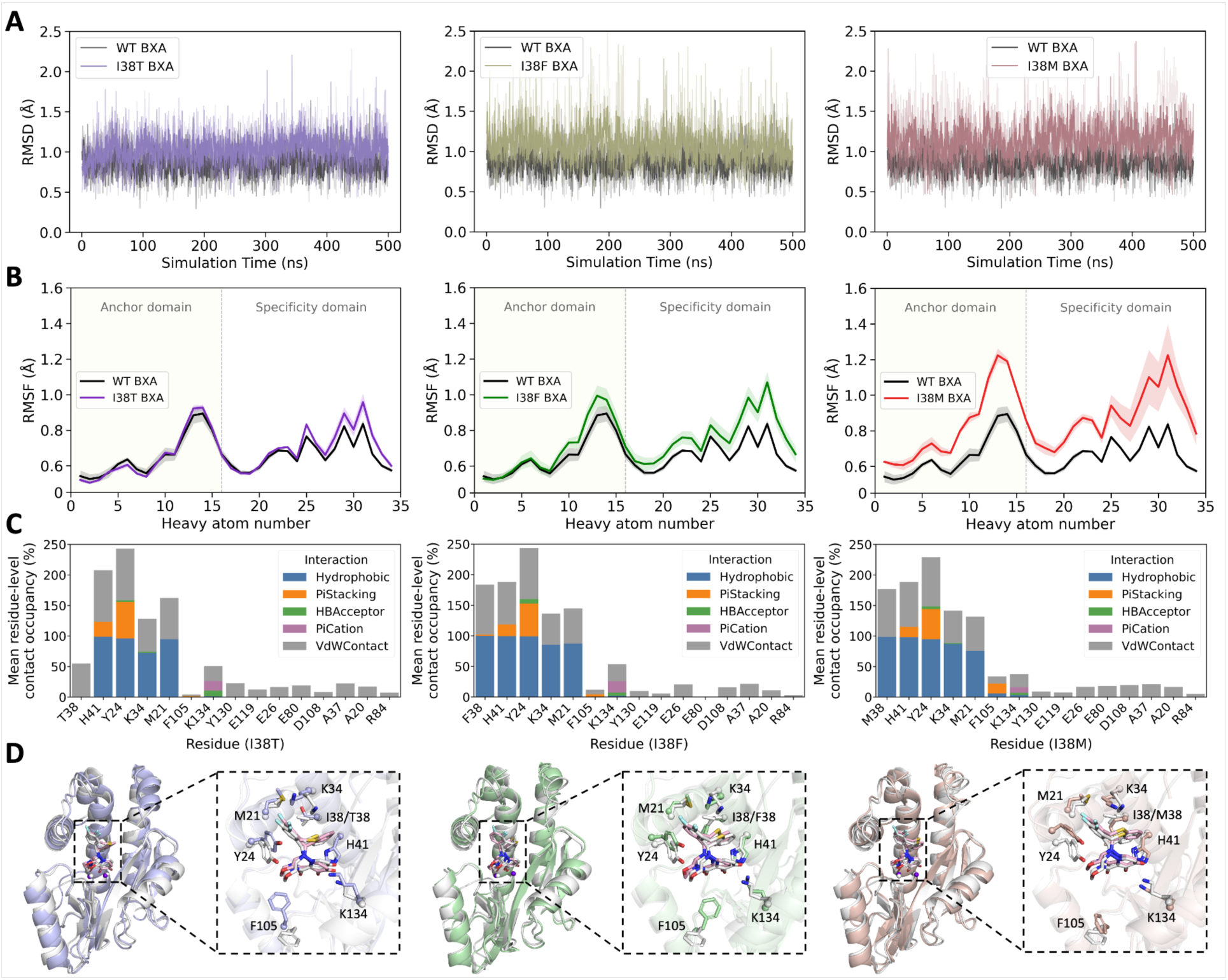
Summary of BXA binding dynamics and interaction profiles in I38 variants. **(A)** BXA RMSD profiles comparing WT with I38T, I38F, and I38M variants. RMSD was calculated relative to the initial structure of each corresponding system. Solid lines represent the average of three independent replicas, while individual replicas are shown as semi-transparent lines with distinct transparency levels. **(B)** Atom-level RMSF profiles of non-hydrogen BXA atoms comparing WT with I38 variants. The BXA anchor domain is highlighted in pale yellow to distinguish it from the specificity domain. **(C)** Residue-level BXA interaction occupancy across the I38 variant trajectories. **(D)** Representative BXA-bound structures of the I38 variants overlaid with WT, shown in gray, highlighting the BXA binding pose and key interacting residues.

For I38F, larger BXA RMSD fluctuations were consistent with atom-level RMSF analysis, which also showed increased ligand mobility primarily localized at the hydrophobic wing of BXA (**Figure 6A-B**). Contact frequency analysis showed that, unlike I38T which abolished the native Ile38-mediated hydrophobic interaction, substitution of Ile38 with the bulkier phenylalanine retained comparable hydrophobic occupancy to the WT. Concurrently, increases were observed in Met21 hydrophobic contacts and Lys134-associated H-bond/π-cation interactions, while Phe105 π-stacking dropped to negligible levels as I38T (**Figure 6C**). Representative structures provided a clear physical explanation for this interaction redistribution. The substitution introduced a larger aromatic side chain that could not pack into the V-shaped difluoro-dihydro-dibenzothiepine wing as compactly as the WT Ile38. Instead, Phe38 occupied a slightly offset position. To maintain residue 38 contact despite this less optimal hydrophobic packing geometry, the benzene ring from the specificity domain shifted inward and downward while the difluorinated aromatic end tilted upward toward Met21, thereby increasing the probability of Met21 contacts. The accompanying adjustment of the anchor domain also likely disrupted the geometric alignment required for Phe105 π-stacking, explaining the loss of this interaction (**Figure 6D**). This less optimal packing geometry was further destabilized by increased Phe38 side-chain dynamics. In contrast to the compact WT Ile38 side chain, the Phe38 aromatic ring underwent repeated conformational reorientations, driving an approximately 0.4 Å increase in side-chain RMSF despite maintaining stable backbone dynamics **(Movie S1)**. Consequently, although residue 38-BXA hydrophobic occupancy remained high, different Phe38 ring atoms alternately engaged BXA, producing a dynamically exchanged contact network rather than a persistently locked hydrophobic interface. Accordingly, residue-level dynamic cross-correlation matrices (DCCM) showed that Phe38-BXA coupling decreased from 0.38 in WT to 0.26 in I38F, indicating weakened coordinated motion despite preserved contact frequency. This loss of coherent packing allowed the BXA hydrophobic wing to sample multiple local micro-poses within the pocket, explaining the increased ligand RMSF. Thus, I38F likely reduces BXA efficacy by converting the native Ile38-centered hydrophobic anchor into a dynamic, exchangeable packing interface that destabilizes the high-affinity WT-like binding geometry.

The I38M system displayed the more pronounced RMSD fluctuations of the Ile38 variants (**Figure 6A**). Atom-level RMSF showing increased ligand mobility was no longer confined to the hydrophobic wing. Instead, it propagated into the metal-chelating anchor domain, driving coupled fluctuations of the catalytic Mn^2+^ ions **(Figure 6B and Figure S6**). Contact frequency analysis showed that the hydrophobic occupancy between Met38 and BXA experienced only a negligible decrease. Concurrently, modest reductions were also observed in Met21 and His41 hydrophobic/aromatic interactions, whereas Lys34 contacts increased slightly **(Figure 6C)**. Representative structures provided a clear spatial rationale for these altered interaction profiles. Similar to I38F, the elongated methionine side chain was unable to pack into the native position, extending downward instead. Consequently, BXA deviated from its WT binding conformation, with the specificity domain shifting upward and inward, leading to less optimal hydrophobic packing with residue 38. This rearrangement was accompanied by outward displacement of the α2 C-terminal region, consistent with the reduced Tyr24 interactions, while simultaneously shifting the specificity wing toward Lys34 and away from His41, explaining the increased Lys34 contact probability and decreased His41 interactions **(Figure 6D)**. Unlike the rigid, β-branched WT isoleucine, the linear aliphatic chain of methionine possesses high rotational degrees of freedom, resulting in an increase of nearly 1 Å in residue 38 side-chain RMSF relative to WT despite similar backbone position. Trajectory inspection further indicated that Met38 was more mobile than Phe38. The linear methionine side chain bent and rotated to maintain hydrophobic contacts with BXA through different atoms, but it failed to impose the stable geometric constraint provided by WT Ile38 **(Movie S2)**. Consistent with this interpretation, residue 38-BXA cross-correlation decreased more strongly in I38M, reaching approximately 0.16 compared to 0.38 in WT, confirming a substantial loss of coherent coupling. Taken together, these findings suggest that I38M compromises BXA efficacy primarily by replacing the compact Ile38 hydrophobic anchor with a highly flexible methionine side chain that fails to maintain a stable and geometrically optimized packing interface. This loss of coherent hydrophobic anchoring destabilizes the global BXA framework and perturbs the apparent positional stability of the Mn^2+^ ions. In an unrestrained physiological environment, substantial BXA fluctuations that displace the anchor domain away from the Mn^2+^ ions may facilitate water-mediated competition for Mn^2+^ coordination while weakening ligand chelation.

### A36V destabilizes the Apo PA_N_ and promotes open pocket conformations

Unlike active-site mutations such as those at Ile38, A36V is located distal to the binding pocket and does not directly interact with BXA. Throughout the simulations in the apo state, A36V exhibited larger loop-excluded backbone RMSD fluctuations than WT, along with a broader replica-averaged distribution, indicating expanded sampling of core conformations and greater replica-to-replica variability. RMSF analysis localizes the increased flexibility primarily to the N-terminal region (residues 1-40) and the C-terminal tail **(Figure 7A-B)**. While PCA calculated from the whole protein showed only a single basin largely overlapping with WT, loop-excluded PCA resolved a broader apo A36V conformational landscape comprising three clusters **(Figure 7C)**. All simulation replicas sampled each cluster but with distinct population preferences, supporting genuine conformational heterogeneity rather than replica-specific sampling artifacts (**Figure S7**). A comparison of the representative structures from each cluster revealed that the principal conformational divergence is localized within the same regions as those with increased RMSF, characterized by local α-helix unwinding. Secondary-structure occupancy calculated with cpptraj quantitatively supported this observation. In addition to reduced α-helical occupancy at the N- and C-terminal ends, residues 22-24 near the α2 C-terminal end showed marked decreases from 0.985, 0.921, and 0.700 in WT to 0.647, 0.456, and 0.312 in A36V, respectively. Reduction was also observed near the beginning of α3, where residues 32-35 decreased from nearly complete α-helical occupancy in WT to 0.910-0.930 in A36V. Residues 36-37 showed the strongest loss, decreasing from 1.00 to 0.34, while residue 38 showed a smaller reduction from 1.00 to 0.86. These results indicate that A36V promotes localized secondary-structure unfolding at both the flexible termini and pocket-adjacent α-helical regions.

**Figure 7.**
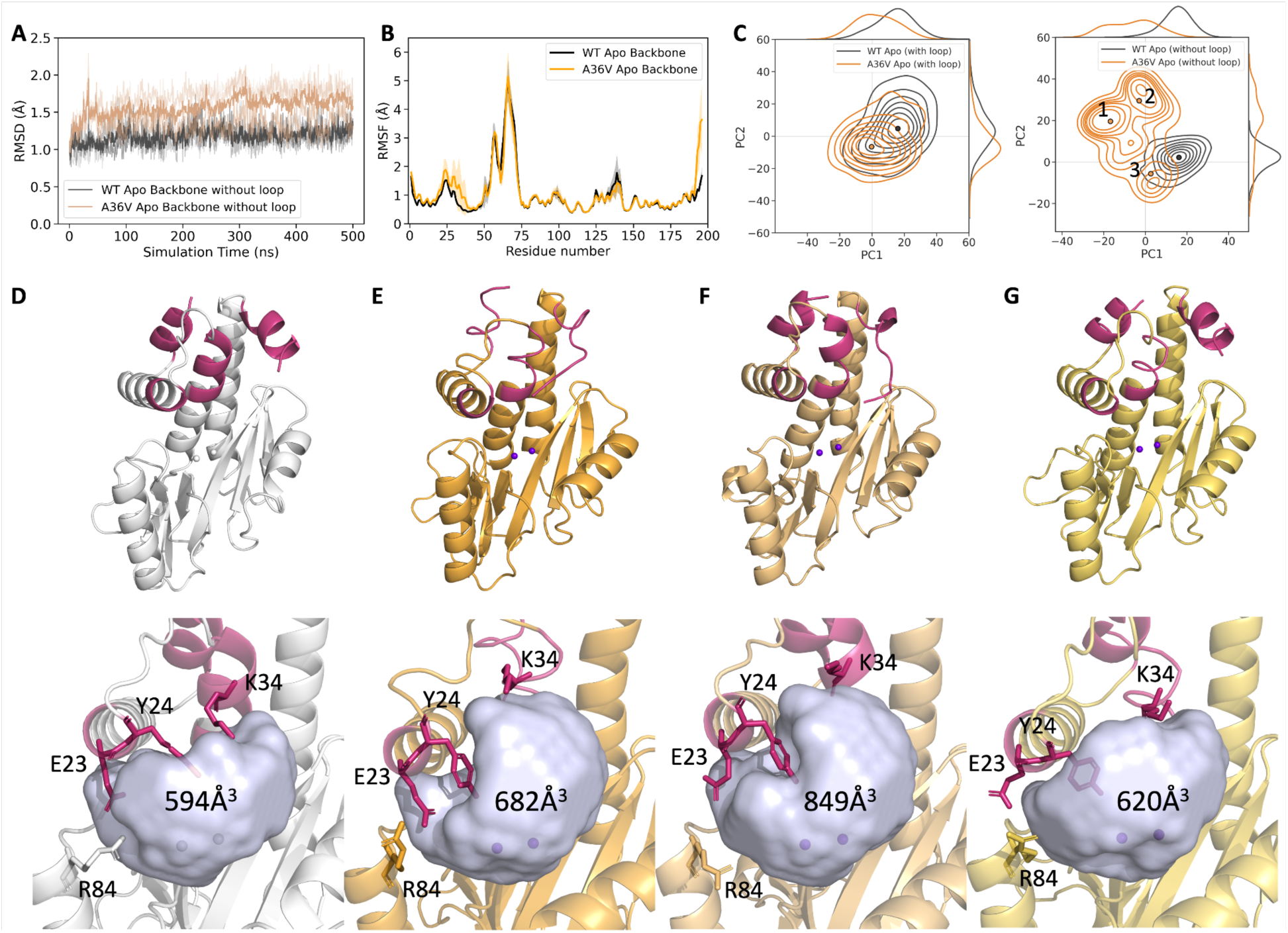
Apo-state simulations reveal local destabilization and altered pocket conformations in A36V PA_N_. **(A)** Loop-excluded backbone RMSD profiles comparing WT and A36V apo systems. **(B)** Residue-level RMSF profiles of WT and A36V apo systems. **(C)** KDE contour plots comparing WT and A36V conformational distributions from full-backbone and loop-excluded PCA projections. Points indicate the representative frames closest to the mean-shift cluster centers for each system in the corresponding PCA space. **(D-G)** Representative structures identified from mean-shift clusters in the loop-excluded PCA space, with corresponding pocket volumes shown in light blue and labeled in Å^3^. WT is shown in D, and A36V structures from clusters 1, 2, and 3 are shown in E-G, respectively. Warm pink highlights regions with reduced α-helical occupancy, and sticks indicate residues contributing to the observed pocket differences among the structures.

To assess whether the observed unwinding of α-helical regions altered pocket geometry, we calculated pocket volumes from representative structures of each basin. In representatives from basins 1 and 2, partial unfolding of the C-terminal segment caused insertion toward the pocket region **(Figure 7E-F)**. However, in the native polymerase context, this segment connects to the PA C-terminal domain and does not intrinsically contribute to the formation of the PA_N_ catalytic pocket, thus the observed intrusion was considered an artifact and was excluded from the pocket volume calculation. Under this definition, representative structures from basins 1 and 2 (located distal to the WT basin center) exhibited expanded pocket volumes relative to WT (682 and 849 Å^3^ versus 594 Å^3^) **(Figure 7D-F)**. In contrast, the representative structure from basin 3 (proximal to the WT center) showed a comparable pocket volume to WT of 620 Å^3^ **(Figure 7G)**. This expansion primarily resulted from local unwinding and conformational rearrangement of the 32-34 region on the α3 helix, which opened additional space adjacent to the pocket. The larger volume in basin 2 was further associated with upward displacement of the α2 helix, widening the pocket boundary. Together, these findings suggest that the broadened apo A36V landscape includes more open pocket conformations, linking localized secondary-structure remodeling of pocket-adjacent regions to altered apo pocket preorganization.

In the BXA-bound simulations, however, overall protein dynamics were highly similar to WT. Backbone RMSF showed only a modest increase in the flexible loop, while the structured core remained comparable to WT, consistent with the PCA showing a broader distribution when calculated with the loop included **(Figure S8A-C)**. BXA maintained a stable binding pose throughout the simulations, with RMSD and RMSF profiles closely matching WT **(Figure S8A-B)**, indicating no substantial destabilization of the bound inhibitor. Contact analysis further supported this conclusion. The overall BXA interaction network in A36V was largely preserved, with only minor local redistribution. Tyr24 π-stacking occupancy decreased from 56.5% to 46.3%, consistent with outward displacement of the α2 C-terminal region, whereas Lys134 H-bond and π-cation contacts modestly increased to 13%, likely reflecting upward displacement of the α5 helix (**Figure S8D-E**). These changes indicate subtle shifts in local contact preference rather than pronounced ligand destabilization.

Previous experimental study^29^ showed that A36V substantially reduces apo PA_N_ stability, with a melting temperature nearly 10 °C lower than WT and insufficient stability for reliable Kd measurement. In contrast, BXA-bound A36V produced a WT-like thermal shift, and its crystal structure showed no notable change in BXA binding conformation. Consistent with these observations, our simulations indicate that A36V does not directly destabilize the bound inhibitor. Instead, A36V primarily broadens the apo conformational landscape and enriches more open pocket states, suggesting that its resistance mechanism is driven by local apo-state destabilization and altered pre-binding conformational sampling rather than unfavorable properties of the bound conformation.

### E23K rewires the α2 electrostatic network and perturbs BXA specificity-wing packing

Residue 23 is located on the α2 helix and, similar to A36V, does not directly interact with BXA when bound. In the apo state, the structural dynamics of E23K remain broadly WT-like, exhibiting no obvious conformational ensemble shift **(Figure S9)**. Instead, the primary dynamic alterations of E23K emerged in the BXA-bound state. Without the flexible loop, the backbone RMSD remained comparable to WT, indicating that the structured core was overall stable in the bound state. However, when the loop was included, RMSD and RMSF showed large replica-to-replica variation in the flexible loop region and higher average fluctuation than WT (**Figure 8A-B**). In contrast to A36V, for which the largest conformational differences were revealed by loop-excluded PCA, E23K displayed broader conformational sampling only when the flexible loop was included in the PCA calculation. In the full-backbone PCA space, E23K separated into three clusters (Figure 8D), whereas the loop-excluded PCA projection showed conformations occupying a single dominant basin. Of the three loop-associated clusters, Cluster 1 represents the dominant state (66.6% of all frames), overlapping with the WT basin, with roughly equal contributions from the second and third MD replicas. Cluster 2 (27.0%) was exclusively sampled by the first MD replica, whereas Cluster 3 (6.5%) was sampled in the first and third replicas. Representative structures from the three clusters further showed that their primary structural differences were localized to the flexible loop **(Figure 8E)**. Notably, similar regions of PCA space are sparsely sampled by other systems, but their populations are too low to form a distinct density mode **(Figure S10)**. This suggests that these loop states are likely inherently accessible within the PA_N_ conformational ensemble, but are significantly enriched by the E23K mutation. While the previous study^66^ has demonstrated that this flexible loop can modulate viral replication by adopting different interaction modes with PB1 and PB2 to stabilize the replicase polymerase, our study utilizes an isolated PA_N_ model lacking the full RdRp context. Therefore, this loop-associated redistribution is strictly reported as an observed conformational feature, without further interpreting it as a definitive functional mechanism.

**Figure 8.**
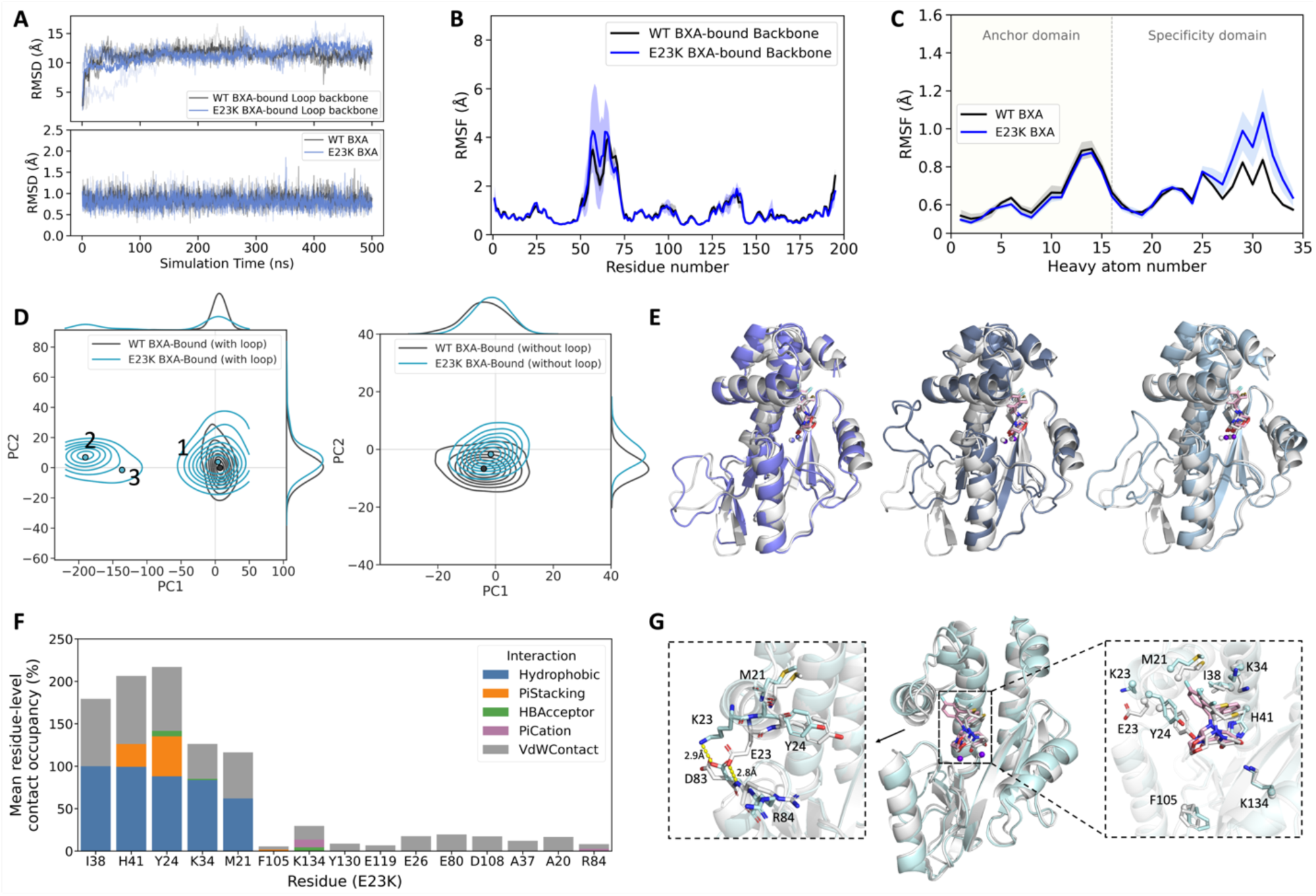
Summary of loop dynamics and BXA binding interactions in the E23K BXA-bound system. **(A)** Loop backbone and non-hydrogen BXA RMSD profiles comparing BXA-bound WT and E23K systems, calculated relative to the initial structure of each corresponding system. **(B)** Residue-level RMSF profiles comparing WT and E23K BXA-bound systems. **(C)** Atom-level RMSF profiles of non-hydrogen BXA atoms bound to WT and E23K systems. **(D)** KDE contour plots comparing WT and E23K PCA projections calculated from the full protein backbone and loop-excluded backbone. Points indicate representative frames closest to the mean-shift cluster centers for each system in the corresponding PCA space. **(E)** Representative structures from the three loop-associated clusters identified in the full-backbone PCA space for the BXA-bound E23K system, shown in the order of clusters 1, 2, and 3 and overlaid with the WT representative structure in gray. **(F)** Residue-level BXA interaction occupancy across the BXA-bound E23K trajectories. **(G)** Representative structures from WT and E23K loop-excluded PCA clusters, with WT shown in gray, highlighting the altered E23K interaction network and key BXA-interacting residues.

Although the overall ligand RMSD of E23K closely resembles that of the WT, atom-level BXA RMSF reveals increased fluctuations in the fluorinated aromatic ring of BXA, indicating a reduced local stability of the inhibitor (**Figure 8A, C**). Contact-frequency analysis further supported this localized perturbation (**Figure 8F**). In E23K, several interactions stabilizing the BXA specificity wing were weakened: hydrophobic contacts with Met21 and Tyr24 decreased from 77.6% and 97.5% in WT to 62.0% and 88.1%, respectively, while Tyr24 π-stacking decreased from 56.5% to 47.2%. At the same time, the π-stacking interaction between Phe105 and the heteroaromatic ring of the anchor domain was nearly abolished, decreasing from 13.9% to almost undetectable levels. Intra-protein interaction analysis suggests that these effects originate from electrostatic rewiring near residue 23. In WT, Glu23 forms a persistent interaction network with Arg84 through both hydrogen-bonding and anionic interactions, with occupancies of 73.8% and 57.9%, respectively. Substitution of negatively charged glutamate with positively charged lysine abolished the native Glu23-Arg84 interaction and instead promoted formation of new Lys23-Asp83 cationic and hydrogen-bonding interactions, with occupancies of 77.0% and 64.2%, respectively. Representative structures further elucidated that these newly formed interactions pull the α2 C-terminal region toward Asp83, shifting residues 20-24 slightly away from the active site. Consistent with this rearrangement, the difluoro-dihydro-dibenzothiepine wing of BXA exhibited a corresponding positional displacement accompanied by a slight retreat of the heteroaromatic ring from the BXA anchor domain, in agreement with the near-complete loss of Phe105 π-stacking interactions (**Figure 8G**).

Collectively, E23K replaces the native Glu23-Arg84 interaction with a new Lys23-Asp83 contact, reshaping the local electrostatic network around the α2 C-terminal region. This rearrangement shifts the pocket-adjacent α2 segment and weakens key Met21-, Tyr24-, and Phe105-mediated hydrophobic/aromatic contacts with BXA. Thus, E23K likely reduces BXA susceptibility through localized electrostatic rewiring and pocket remodeling that destabilize the inhibitor interaction network.

## Conclusions

In this study, we first benchmarked multiple metal-modeling strategies in apo and BXA-bound WT PA_N_ to establish a reliable framework for describing the challenging bimetallic Mn^2+^ active center. Based on this validated framework, we performed MD simulations of WT and five clinically relevant variants, with three independent 500 ns replicas for each system, resulting in 1.5 μs of sampling per variant. These simulations revealed diverse dynamic mechanisms underlying reduced BXA susceptibility. Consistent with previous studies, I38T reduced BXA binding stability primarily by disrupting the native Ile38-mediated hydrophobic anchoring of the BXA specificity wing. In contrast, I38F and I38M retained apparent residue 38-BXA hydrophobic contact, but their larger and more flexible side chains failed to reproduce the compact and coherent WT-like packing geometry required to immobilize the BXA. Distinctly, A36V reconfigures the apo-state conformational landscape. Mutation-induced destabilization of the local α-helix region enriches more open pocket conformations, thereby perturbing the pre-binding organization. Furthermore, E23K reshapes the binding environment through localized electrostatic rewiring. The replacement of the native Glu23-Arg84 interaction with a novel Lys23-Asp83 network shifts the α2-terminal region and weakens key hydrophobic and aromatic contacts involving Met21 and Tyr24 that normally stabilize the BXA specificity wing. Collectively, this work not only provides methodological guidance for modeling complex bimetallic active sites, but also establishes a unified dynamic framework for understanding BXA resistance. By revealing how distinct resistance mutations reshape PA_N_ conformational ensembles and BXA binding modes, these findings offer a structural and mechanistic basis for future design and optimization of next-generation influenza endonuclease inhibitors.

## Data Availability

All input structures and representative structures used for analysis can be found at https://github.com/SztainLab/PAN_parameters available upon publication.

## Code Availability

Simulation input scripts can be found at https://github.com/SztainLab/PAN_parameters available upon publication. Amber is available at: https://ambermd.org/.

## Acknowledgements

We thank Prof. Kenneth M Merz Jr., Dr. Zhen Li, and Subhamoy Bhowmik for helpful discussions and for providing C_4_ terms between Mn^2+^ and hydroxide oxygen, and between hydroxide oxygen and TIP3P water. This material was made possible, in part, by a Cooperative Agreement from the United States Department of Agriculture’s Animal and Plant Health Inspection Service (APHIS) award AP26VSSP0000C021 to TS. It may not necessarily express APHIS’ views. This work was partially supported by the start-up funds from Loyola University Chicago (to PL).

## Author Contributions

Conceptualization: TS, Supervision: TS and PL, experiments: LW, analysis: LW, with contribution from all authors, manuscript preparation: LW, with contribution from all authors.

