## Supplementary Information for "Parameterization of the PA endonuclease bimetallic center reveals the dynamics of clinically relevant mutations"

### Table of Contents

### Supplementary Figures

**Figure S1.** Bimetallic coordination geometry of apo $PA_N$ using the 12-6-4 nonbonded model.
**Figure S2.** Bimetallic coordination geometry of apo $PA_N$ using the bonded model.
**Figure S3.** Bimetallic coordination geometry of apo $PA_N$ using the hybrid model.
**Figure S4.** Apo $PA_N$ dynamics and conformational sampling of I38 variants.
**Figure S5.** BXA-bound $PA_N$ dynamics and conformational sampling of I38 variants.
**Figure S6.** Distribution of catalytic $Mn^{2+}$ RMSF values across BXA-bound WT and mutant $PA_N$ systems.
**Figure S7.** Apo A36V cluster assignments shown by individual replicas.
**Figure S8.** Dynamics and interaction profiles of BXA-bound A36V $PA_N$.
**Figure S9.** Apo $PA_N$ dynamics and conformational sampling of the E23K variant.
**Figure S10.** Full-backbone PCA projection comparing conformational sampling across all BXA-bound $PA_N$ systems.

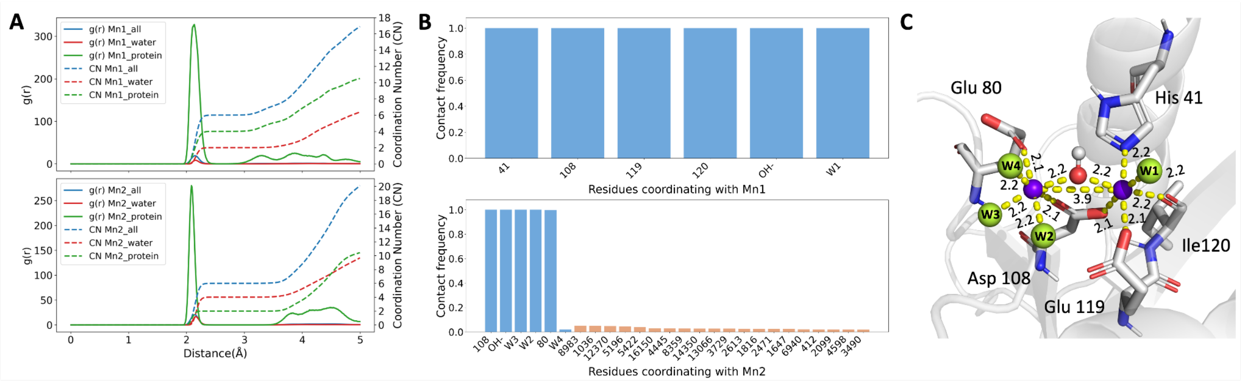


**Figure S1. Bimetallic coordination geometry of apo PA$_N$ using the 12-6-4 nonbonded model. (A)** RDFs were calculated separately for each $Mn^{2+}$ ion. Integration of the first coordination shell yielded coordination numbers consistent with the expected PA$_N$ metal environment. **(B)** Contact-frequency analysis of the first-shell coordinating ligands of Mn1 and Mn2. For Mn1, the blue bars indicate that the native coordinating ligands remain stably coordinated throughout the simulation. For Mn2, the native ligands show the same near-complete occupancy, whereas the W4 site is maintained by dynamically exchanging solvent waters. The wheat-colored bars represent the top 20 replacement waters among 159 solvent waters that transiently occupy this site. Although each individual replacement water has low occupancy, their cumulative contact frequency is also nearly 1. **(C)** Representative structure showing the preserved $Mn^{2+}$ first-shell coordination geometry under the 12-6-4 model. Average coordination distances are labeled in Å.

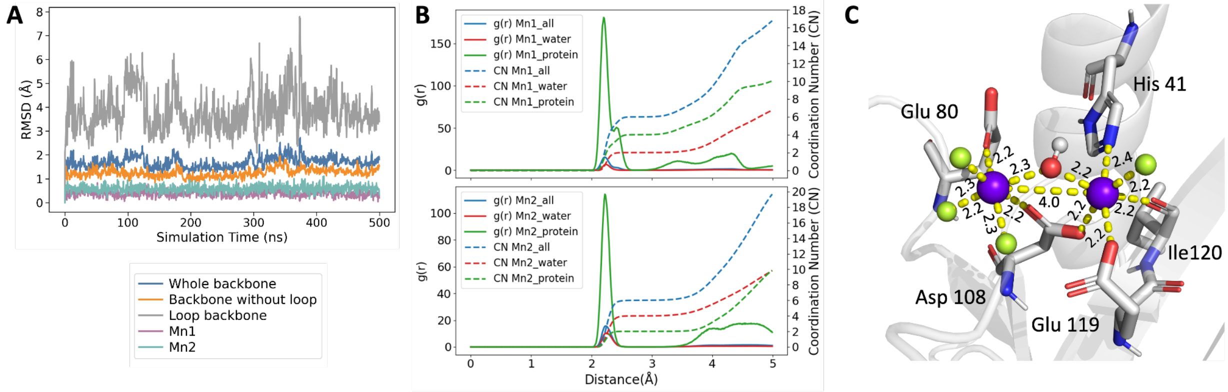


**Figure S2. Bimetallic coordination geometry of apo PA$_N$ using the bonded model. (A)** RMSD profiles of the apo PA$_N$ system. **(B)** Average RDFs and coordination numbers for the two $Mn^{2+}$ ions. **(C)** Representative structure of the bimetallic active center, with average coordination distances labeled in Å.

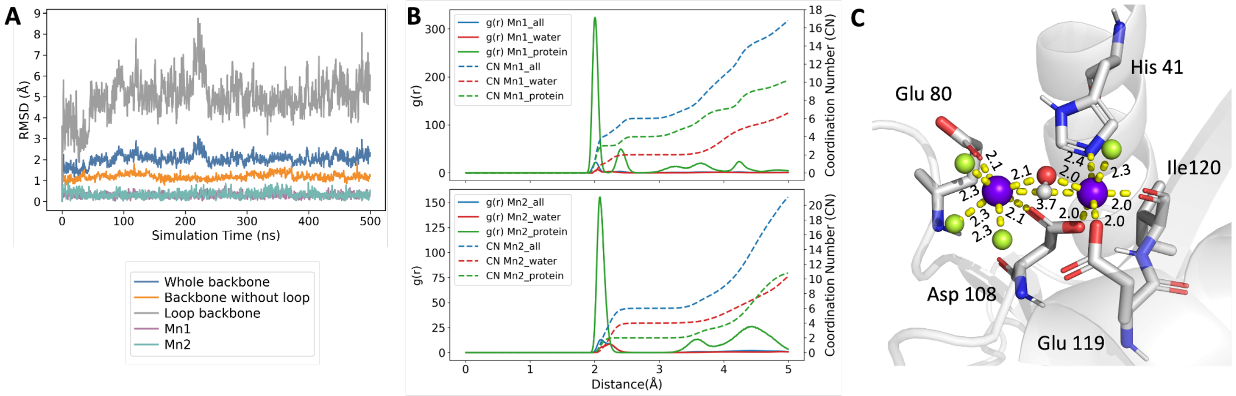

**Figure S3. Bimetallic coordination geometry of apo $PA_N$ using the hybrid model. (A)** RMSD profiles of the apo $PA_N$ system. **(B)** Average RDFs and coordination numbers for the two $Mn^{2+}$ ions. **(C)** Representative structure of the bimetallic active center, with average coordination distances labeled in Å.

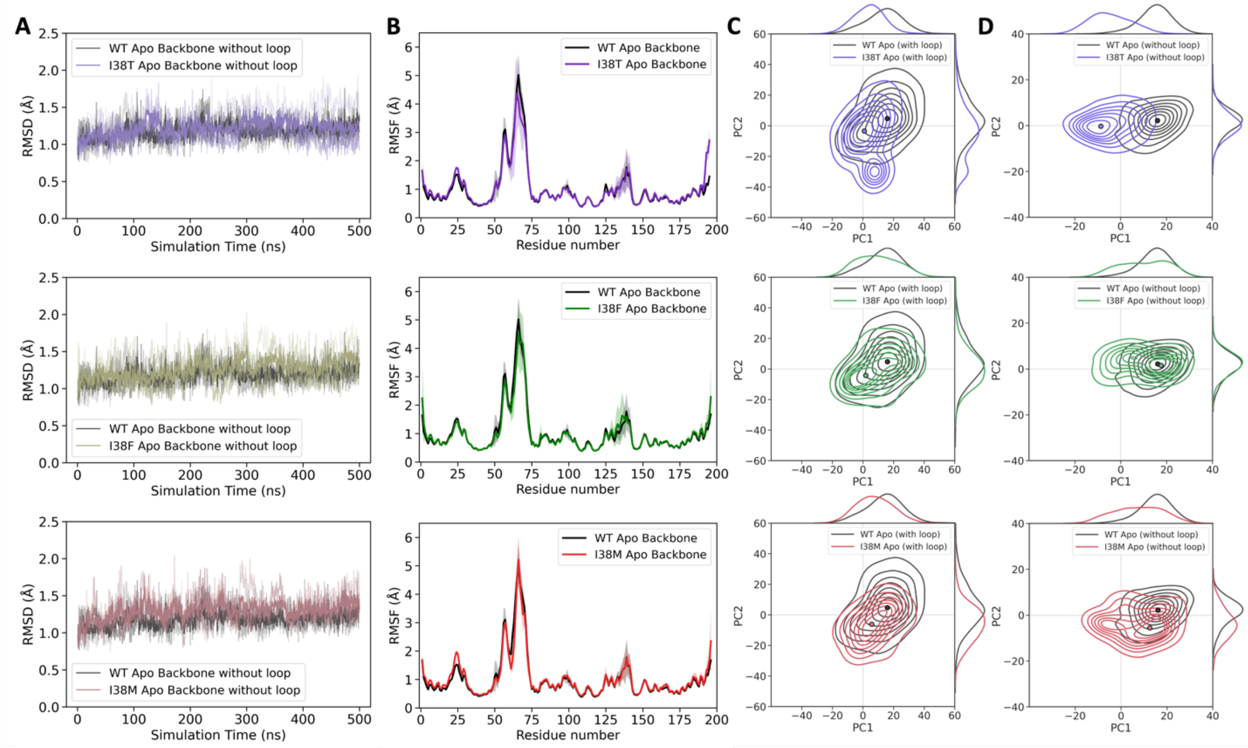

**Figure S4. Apo $PA_N$ dynamics and conformational sampling of I38 variants. (A)** Backbone RMSD profiles excluding the flexible loop for WT and I38T, I38F, or I38M apo systems. RMSD was calculated relative to the initial structure of each corresponding system. Solid lines represent the average of three independent replicas, while individual replicas are shown as semi-transparent lines with distinct transparency levels. **(B)** Residue-level RMSF profiles comparing WT with each apo I38 variant. Shaded regions indicate standard deviations across three independent replicas. **(C)** KDE contour plots of PCA projections for apo WT and I38 variants calculated from the full protein backbone, with marginal KDE curves showing the corresponding PC1 and PC2 distributions. Points indicate the representative frames closest to the mean-shift cluster centers for each system in the corresponding PCA space. **(D)** Corresponding KDE contour plots for apo WT and I38 variants calculated from backbone PCA projections after excluding the flexible loop.

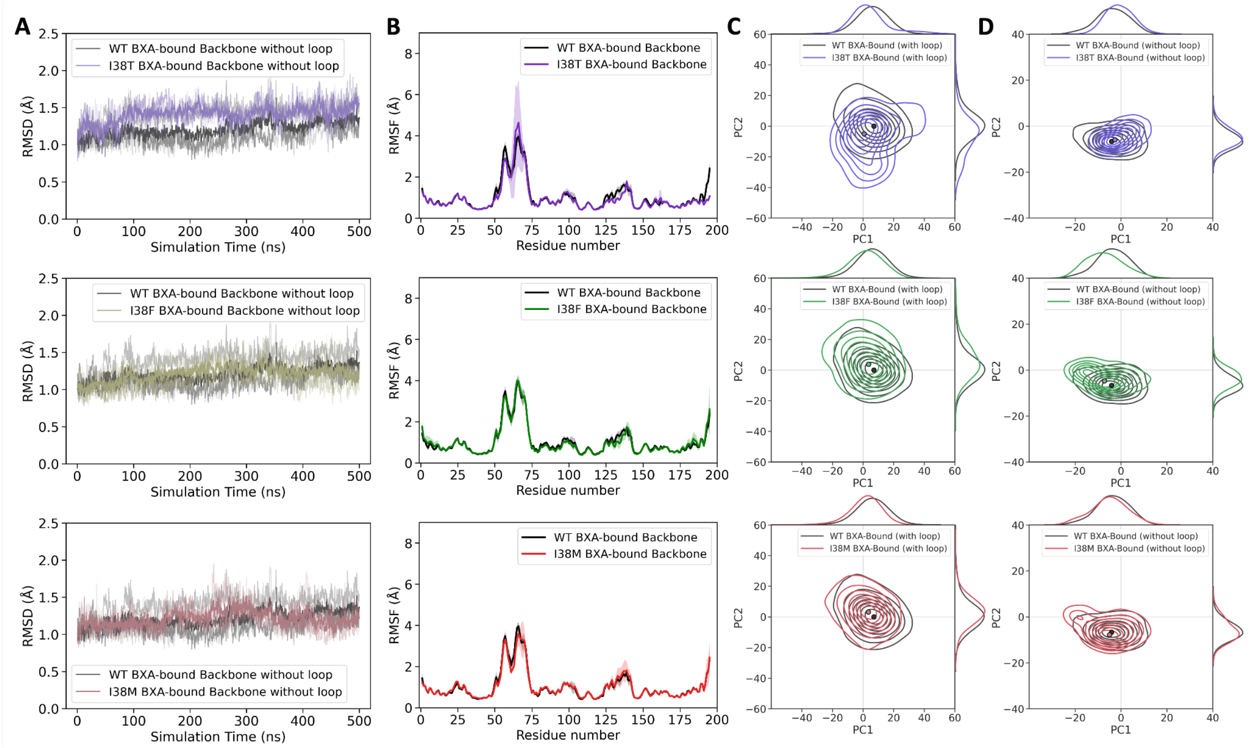


**Figure S5. BXA-bound $PA_N$ dynamics and conformational sampling of I38 variants. (A)** Backbone RMSD profiles excluding the flexible loop for WT and I38T, I38F, or I38M BXA-bound systems. **(B)** Residue-level RMSF profiles comparing WT with each BXA-bound I38 variant. Shaded regions indicate standard deviations across three independent replicas. **(C)** KDE contour plots of PCA projections for BXA-bound WT and I38 variants calculated from the full protein backbone, with marginal KDE curves showing the corresponding PC1 and PC2 distributions. **(D)** Corresponding KDE contour plots for BXA-bound WT and I38 variants calculated from backbone PCA projections after excluding the flexible loop.

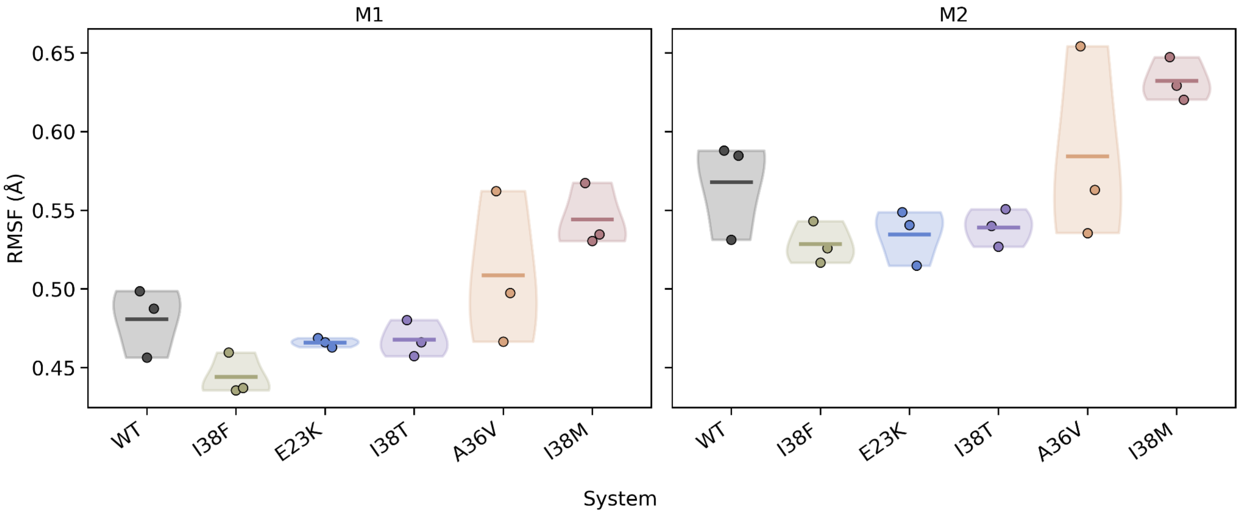


**Figure S6. Distribution of catalytic $Mn^{2+}$ RMSF values across BXA-bound WT and mutant $PA_N$ systems.** Violin plots show M1 and M2 RMSF values calculated from three independent replicas for each BXA-bound system. Points represent individual replica RMSF values.

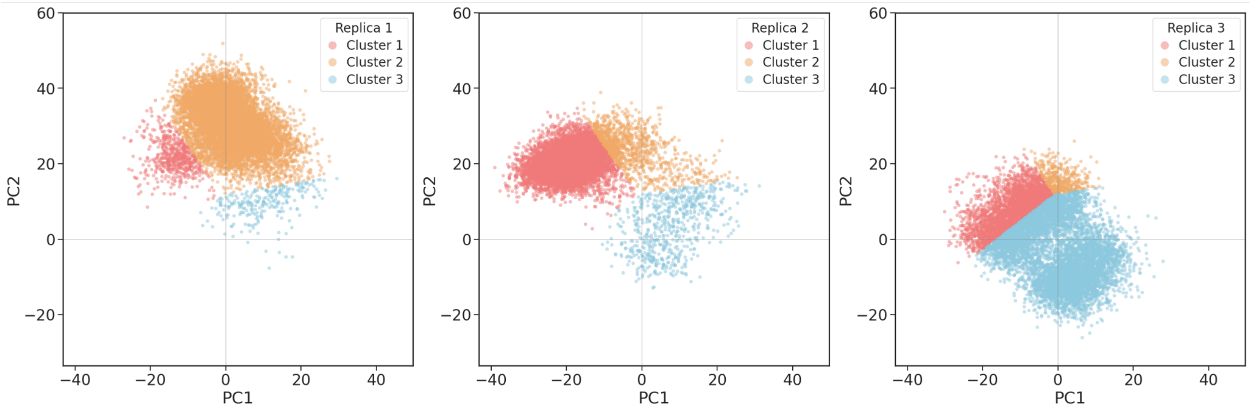


**Figure S7. Apo A36V cluster assignments shown by individual replicas.** Clustering was performed using the combined frames from all replicas, and frames are shown separately by replica with colors indicating cluster assignment.

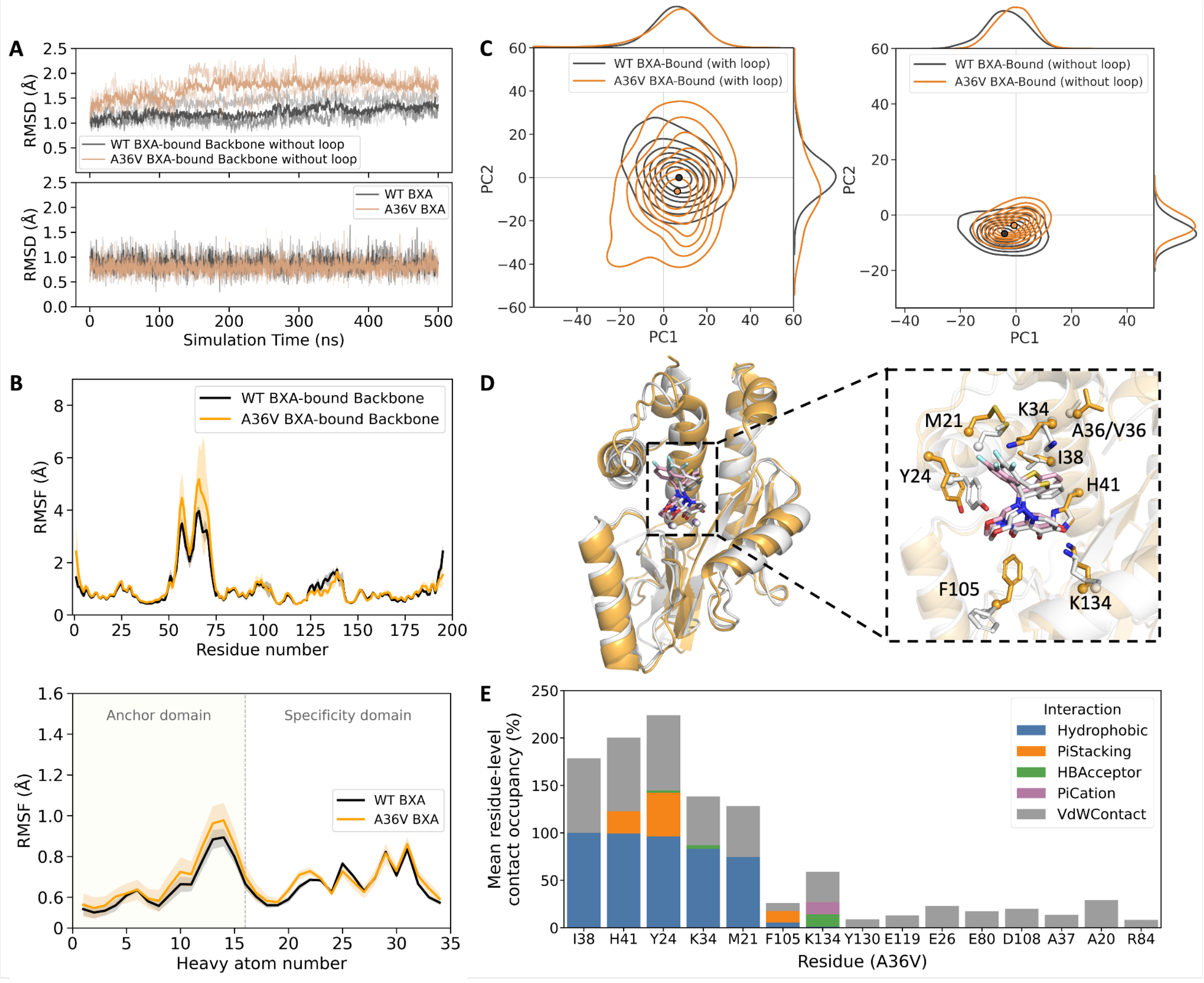


**Figure S8. Dynamics and interaction profiles of BXA-bound A36V $PA_N$. (A)** Loop-excluded backbone and non-hydrogen BXA RMSD profiles comparing BXA-bound WT and A36V systems, calculated relative to the initial structure of each corresponding system. **(B)** Residue-level protein

RMSF profiles and atom-level RMSF profiles of non-hydrogen BXA atoms in WT and A36V BXA-bound systems. **(C)** KDE contour plots comparing WT and A36V PCA projections calculated from the full protein backbone and loop-excluded backbone. **(D)** Representative BXA-bound A36V structure overlaid with WT, shown in gray, highlighting the BXA binding pose and key interacting residues. **(E)** Residue-level BXA interaction occupancy across the BXA-bound A36V trajectories.

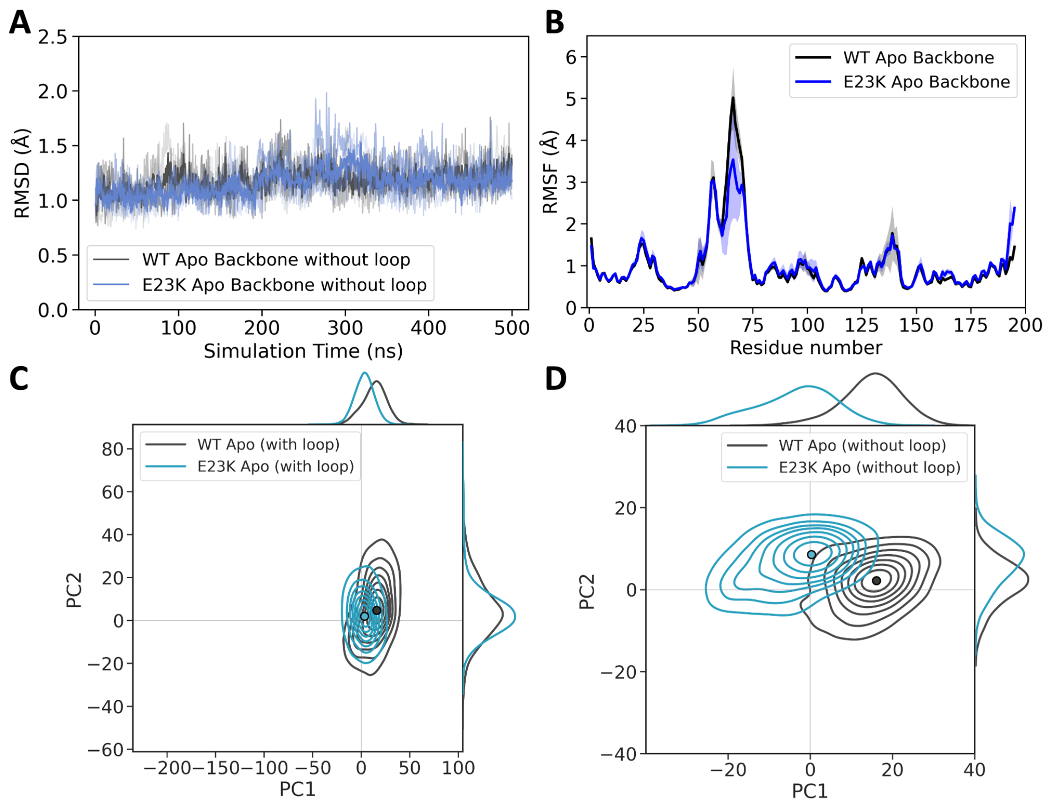


**Figure S9. Apo $PA_N$ dynamics and conformational sampling of the E23K variant. (A)** Loop-excluded backbone RMSD profiles comparing WT and E23K apo systems. **(B)** Residue-level RMSF profiles of WT and E23K apo systems. **(C)** KDE contour plots of PCA projections from WT and E23K apo trajectories calculated using the full protein backbone. **(D)** Corresponding KDE contour plots calculated from loop-excluded backbone PCA projections.

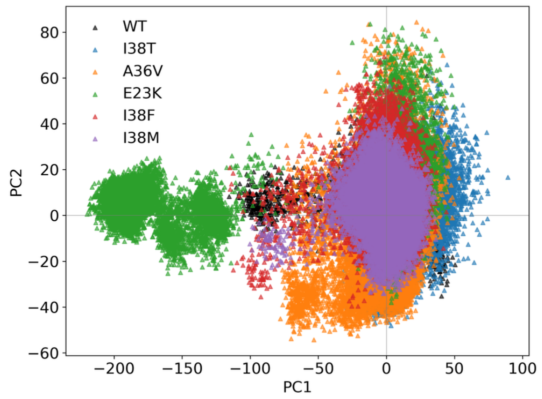


**Figure S10. Full-backbone PCA projection comparing conformational sampling across all BXA-bound $PA_N$ systems**.
